# Domain-informed effector gene assignment exposes training-data dependence of drug target prioritisation from GWAS

**DOI:** 10.64898/2026.11.17.766419

**Authors:** Leonhard Kohleick, Kurt Herzog, Carl Beuchel, Martijn Zoodsma, Claudia Langenberg, Benjamin Wild, Maik Pietzner

**Affiliations:** Health Data Modelling, Berlin Institute of Health at Charité - Universitätsmedizin Berlin, Germany; Computational Medicine, Berlin Institute of Health at Charité - Universitätsmedizin Berlin, Germany; Digital Health, Berlin Institute of Health at Charité - Universitätsmedizin Berlin, Germany; Precision Healthcare University Research Institute, Queen Mary University of London, UK

## Abstract

About 90% of trait-associated lead variants lie outside protein-coding regions, so nominating the effector gene at genome-wide association study (GWAS) loci remains the principal bottleneck in translating associations into biological insight. Since no experimentally resolved ground truth exists at scale, every method rests on quasi-true-positive sets dominated by a single line of evidence. Here, we benchmarked established prioritisation methods across three independent evaluation sets, trained five models each informed by a different biological domain, and annotated lead variants for six common diseases. Performance did not transfer across domains (AUPRC 0.91 to 0.46 for the same method), and we traced this to how differently models weight the distance between variant and gene (p<5.3×10^-123^), not to how they were built. Curation assumptions propagated into predictions and can become self-confirming, such as druggability in the Open Targets-informed model. The five models shared only 17.1% to 31.8% of effector gene assignments, converging on established disease biology but diverging where GWAS are expected to yield novel insight. Recovery of 340 approved drug targets differed markedly between models, and no way of combining them, by averaging scores (38.8%) or training on all curations jointly (36.5%), approached the 52.1% recovered across all five together. Our workflow matched Open Targets L2G across three annotation sets without prior biological knowledge and exceeded it where curated knowledge already linked gene and disease. We provide the Domain-informed EFfector gene Assignment framework (DEFA; https://github.com/lkohleick/DEFA), which reports rather than hides the curation assumption behind each gene assignment so that it can be examined and tested.

## Introduction

Two decades of genome-wide association studies (GWAS) established that disease risk- and quantitative trait-associated variants reside overwhelmingly outside protein-coding regions, with about 90% of published lead variants mapping to the regulatory genome^1,2^. This created a challenge that, despite substantial methodological progress, remains only partially resolved: nominating the putative effector gene among those encoded in proximity, the so-called variant-to-gene (V2G) challenge. Genomic proximity has been a common guide but often misses putative effector genes, since regulatory elements frequently bypass their nearest neighbour^3^ and different classes of genetic signal (e.g., eQTLs versus disease associations) are biased towards different regulatory elements, such that the positional relationship between variant and gene differs systematically^4^. Confidently linking effector genes at non-coding loci has therefore become the major bottleneck in translating GWAS findings into tangible biological insight and therapeutic hypotheses, illustrated by the dependence of clinical success of genetically supported drug candidates on certainty of the causal gene rather than the strength of the association alone^5^.

Despite this centrality, the community has not converged on a standard for generating or reporting effector gene predictions, and independent studies of the same endpoint have nominated different genes even at overlapping loci, agreeing on the top-ranked gene at only 50% to 75% of shared loci across Alzheimer’s disease, heart failure and stroke^6^. How far this discordance extends is largely unknown, because most published lists rank several genes per locus without indicating which is the most likely effector. Scalable frameworks addressing this include MAGMA^7^, PoPS^8^ and FLAMES^9^, which integrate functional and gene-level annotations. In parallel, domain-specific frameworks have been developed that score candidate genes by biological proximity and plausibility^10,11^. Statistical colocalization with QTLs has been widely used for the same purpose^12^, but is restricted to the coverage of disease-relevant cell types and states, which are often unknown and are frequently the very target of hypothesis-generating GWAS^4^.

Since no experimentally resolved ground truth exists for assigning putative effector genes at scale for non-coding loci, existing frameworks are trained and evaluated on a curated set of quasi-true positives, i.e., gene assignments treated as correct in the absence of experimental proof. Each ‘gold standard’ set is further dominated by a single line of evidence. For example, the Open Targets gold standard, used to train L2G^13^, rests on expert- and data-informed annotation, including loci at which the same gene underlies a rare disorder phenotypically similar to the GWAS endpoint. Similarly, the training sets of PoPS and FLAMES rest on the assumption of allelic series. Where statistical fine-mapping identifies both a coding and an independent non-coding signal at the same locus, credible sets containing no coding variant inherit the gene assignment made from the coding variant^8,9^. However, it is currently unclear: 1) whether performance measured on one such set transfers to loci governed by a different regulatory context, 2) how the choice of data curation for model training influences biology inferred from GWAS results, and 3) whether any single curation assumption is sufficient to recover established drug targets.

Here, we show that the choice of building the training set and its inherent assumptions, rather than model architecture, governs both performance and biological interpretation. We find that neither established methods nor five newly built models, each informed by a different biological domain and enriched with prior biological knowledge, performed consistently across quasi-true-positive sets derived from independent lines of evidence. We show that differential model weights for genomic distance (p<5.3×10^-123^) rather than curation quality explained this discrepancy, suggesting domain-specific regulatory mechanisms. Applying every classifier to the same lead variants of six common diseases, we observed convergence of established disease biology but divergence where GWAS are expected to yield novel insights, with only 17.1% to 31.8% of effector genes shared among all five trained classifiers. Finally, recovery of approved drug targets differed markedly between models, and no strategy to combine them approached the 52.1% recovered across all five together; the resulting workflow matched Open Targets L2G without prior biological knowledge and exceeded it by 7.9 to 20.0 percentage points with it. We therefore propose the Domain-informed EFfector gene Assignment framework (DEFA) building on a plurality of domain-specific V2G models whose nominations are combined at the score level and corroborated by orthogonal evidence to advance translation of statistical associations into testable biological hypotheses.

## Results

### Performance of established prioritisation methods does not transfer across biological domains

To first evaluate the performance of state-of-the-art gene prioritisation frameworks, we constructed three distinct evaluation datasets grounded in independent biological reasoning. These datasets leveraged three independent lines of evidence (**Fig. 1a-c; see Methods**): (1) metabolite QTLs (mQTLs) with cross-study validated effector gene assignments (e.g., *BCAT2* and Isoleucine; n=62 genes), (2) GWAS loci that recapitulate known drug – target – indication triplets (e.g., *TSLP* encoding for thymic stromal lymphopoietin, the target of Tezepelumab approved for severe asthma; n=29 genes) that match the indications of FDA-approved therapies, and (3) trans-protein quantitative trait loci (pQTL) that correspond to ligand-receptor pairs or protein complex partners (‘trans-pQTL signals – e.g., *PLAU* and the receptor *PLAUR*; n=101 genes).

**Fig. 1.**
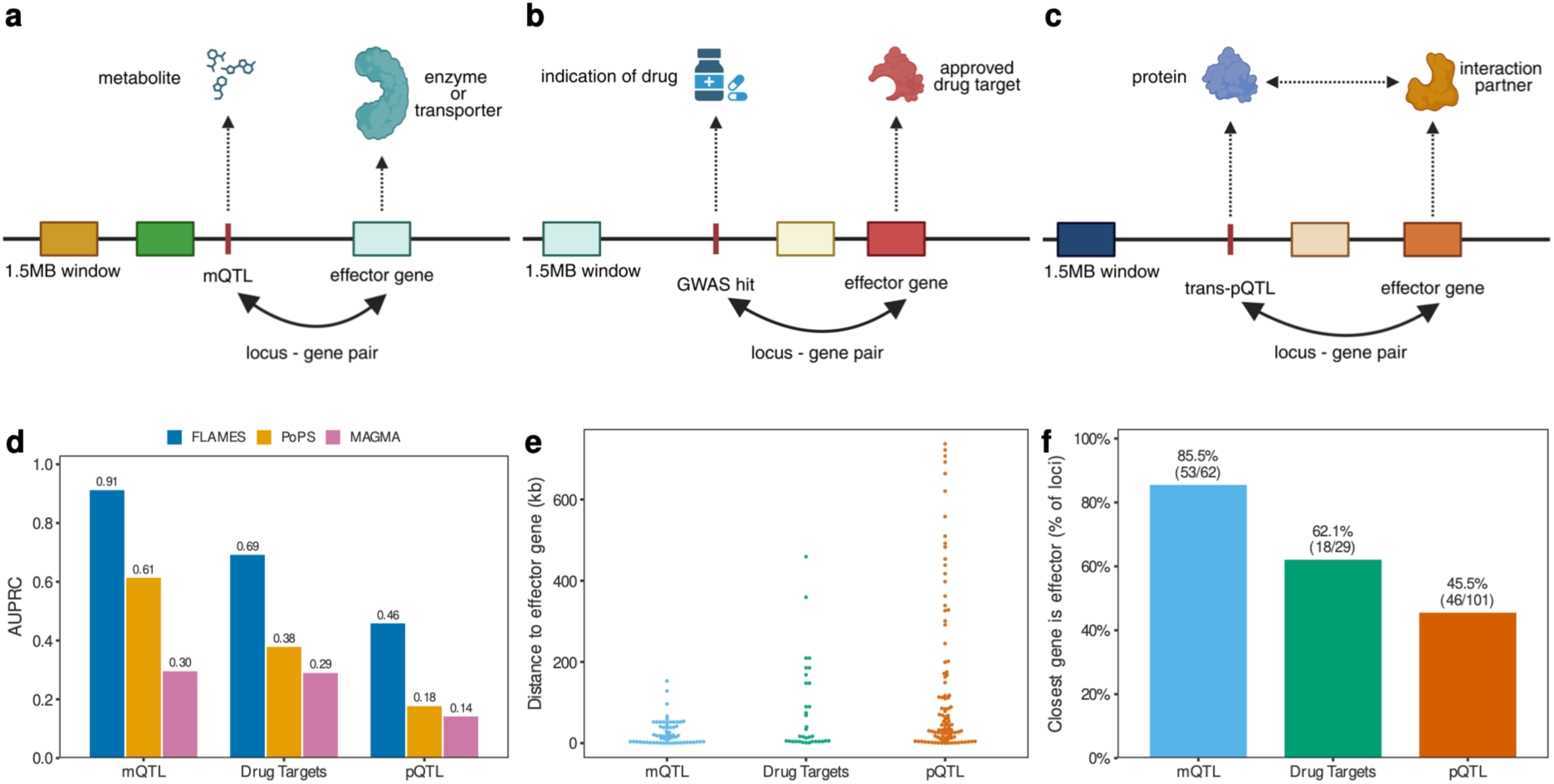
State of the art effector gene methods perform differently across biological domains. **a** We collated mQTL credible sets with high confidence predicted effector genes from two individual studies **b** Harnessing drug-target genetic signals to define “gold-standard” loci. We integrated drug target data from Open Targets^14^ with the GWAS Catalog to identify regional associations within 750 kb of an approved drug-target gene body. **c** Schematic of the *trans*-pQTL-based effector gene assignment. We prioritised candidate effector genes within a 1.5Mb window of the sentinel variant if the gene product participated in a validated ligand-receptor pair or a shared protein complex **d** AUPRC for the three distinct benchmarking datasets (mQTL, Drug Targets, pQTL) **e** Distribution of the distance to effector gene in the datasets **f** percentage when the true effector gene is also the closest gene per dataset. Created in BioRender: https://BioRender.com/k1kkxfb

Across all three datasets, we observed widely varying performance for three state-of-the-art effector gene prioritisation methods, namely MAGMA^7^, PoPS^8^, and FLAMES^9^ (**Fig. 1d and Supplementary Fig. 1; Supplementary Tab. 1)**. Performance dropped as the genomic distance between lead variants and their effector genes grew larger and more variable across datasets (**Fig. 1e,f**). For example, FLAMES performed exceptionally well for mQTLs (area under the precision recall curve (AUPRC): 0.91) but performance dropped strongly for drug target loci (AUPRC: 0.69) and was worst for trans-pQTLs (AUPRC: 0.46) hardly outperforming closest gene annotation (**Fig. 1d**). Similar variation in predictive performance for tools such as PoPS, which do not incorporate genomic proximity in effector gene prediction, suggested that not only local but also global genomic context differs across biological domains, demonstrating the need for training domain-specific effector gene prioritisation pipelines.

### Domain-informed classifiers perform well in-domain but transfer poorly

Having observed poor domain transferability of established effector gene prioritisation methods, we next trained domain-specific effector gene prioritisation models using five distinct quasi-true-positive V2G datasets across a harmonised set of functional genomic features^9^ (**Supplementary Tab. 2**). Each dataset rests on a distinct biological assumption about how variants act on effector genes, defining the domain on which each model was trained and augmenting those reported before: 1) mQTL-informed (489 loci with n=272 unique effector genes), 2) pQTL-informed (416 loci with n=176 unique effector genes), 3) the Open Targets gold-standard set (441 loci with n=272 unique effector genes)^13^, 4) an exonic variant-informed dataset alike FLAMES^9^ (698 loci with n=553 unique effector genes), and 5) a credible-set-informed dataset alike what was used to develop PoPS^8^ (882 loci with n=258 unique effector genes). To incorporate global biological context into our models, we augmented the feature space using metrics calculated on a biomedical knowledge graph^15^ (**Fig. 2a**). Each model showed moderate to good performance on five held-out validation sets (AUPRC: 0.24 – 0.94), but we observed poor cross-domain transferability, in line with earlier findings (Fig.2b). Graph-derived features, i.e., connectivity metrics linking candidate genes to the disease or trait within the knowledge graph, improved performance compared to functional genomic features by on average 16.5% within datasets during cross-validation (range: 11.8%-33.1%) and 12.6% across datasets, with the largest within-dataset gain for the pQTL- and mQTL-informed models, although this gain did not transfer to other datasets (Fig. 2c and Supplementary Tab. 3).

**Fig. 2.**
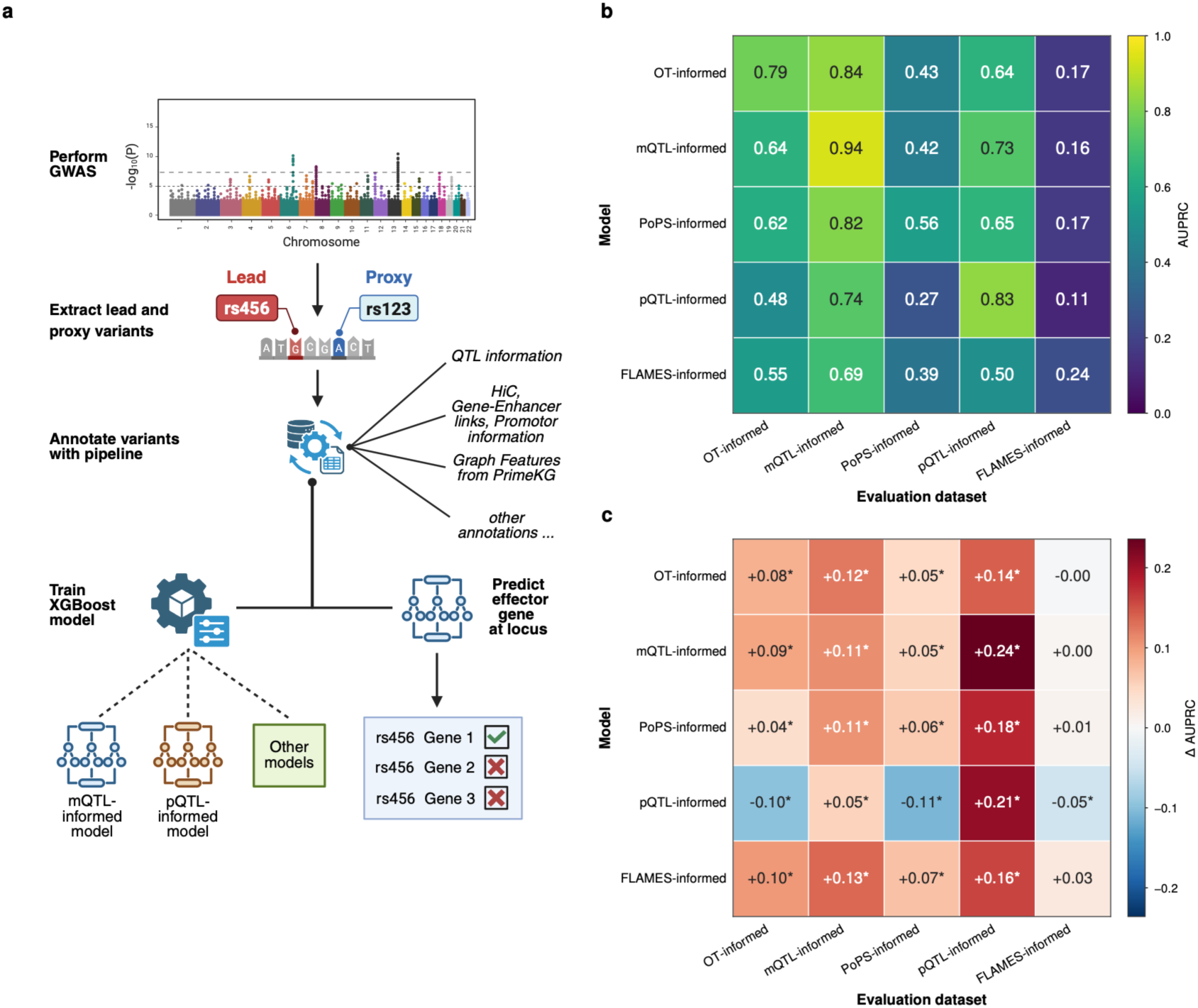
Integrated annotation and training framework for biological domain-specific models. **a**, Schematic of the multidimensional annotation pipeline. Sentinel variants and their respective proxies are annotated by their functional evidence and by their connectivity within the PrimeKG knowledge graph. The framework then either trains XGBoost models on annotated variants or applies previously trained models to a new set of variants. **b**, Comparison of models on a held-out set of variants from each domain reveals marked heterogeneity in performance (AUPRC): no single model out of five performs consistently across all five domains, and inter-model performance varies substantially within domains. **c**, Graph-derived features add the most discriminative power to the pQTL- and mQTL-informed models, and comparatively little to the PoPS- and FLAMES-informed models. Created in BioRender: https://BioRender.com/ueeaxs9

### Differential feature importance across biological domains of effector genes

To understand what drove poor model transferability even though all models were trained on an identical feature set and differed only in their quasi-true-positive labels, we tested each feature in turn for differential associations across domains using logistic regression (Methods). More than half (153 out of 269) of the features showed significantly different (p-value<1.8×10^-4^) associations across the five quasi-true-positive sets (**Fig. 3a**, upper panel, and **Supplementary Tab. 4**), aligning with the poor transferability of model weights beyond the training dataset. Distance to the footprint of the likely causal gene showed the greatest variation in effect sizes (p_interaction_<5.3×10^-123^; **Fig. 3b**), with strongest effects for the mQTL-informed dataset, consistent with metabolite signals acting largely through proximal enzymes and transporters, and least strong effects for the dataset compiled using exonic variant guidance (**Fig. 3b**). Adjusting for a differential association of genomic distance strongly attenuated interaction results for most other features (**Fig. 3a**, lower panel; **Supplementary Tab. 4**). The 29 persistent findings (p-value<1.8×10^-4^) included biologically informed graph-derived features (**Fig. 3c-e**). Graph distance showed stronger effects on pQTL-informed gene sets; druggability was most strongly associated within the Open Targets gold-standard set (odds ratio (OR): 1.93, p-value<3.2×10^-98^) and weakest in the FLAMES-informed dataset (OR: 1.05, p-value=0.22); and measures of evolutionary conservation, e.g., CADD scores, were most strongly associated with positive gene sets derived from mQTLs (OR: 3.12; p-value<1.6×10^-296^).

**Fig. 3.**
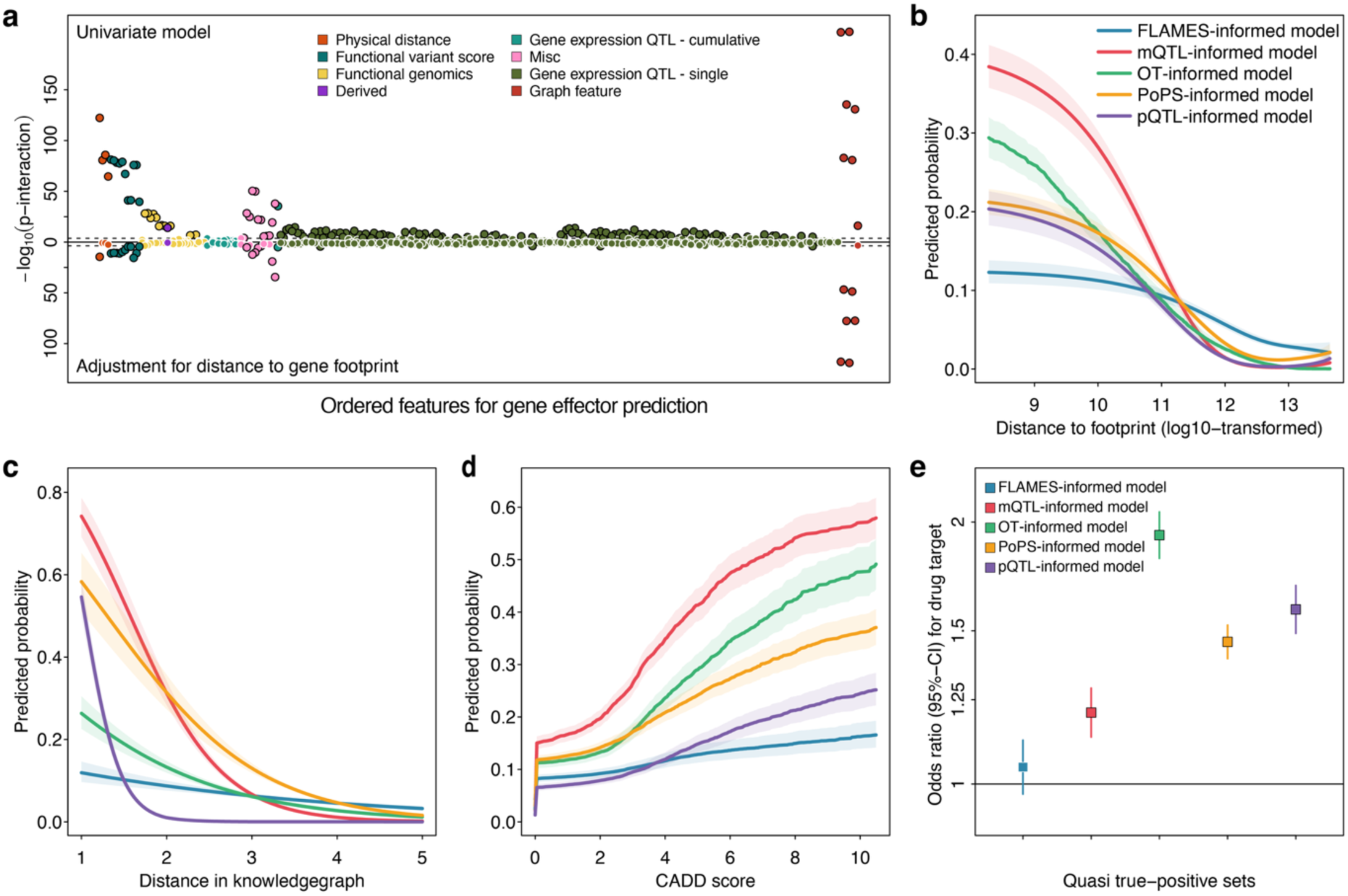
Differential feature importance across quasi-true-positive effector gene sets. **a** Results from systematic interaction testing using logistic regression models on differential association estimates between effector gene prediction features across true positive sets. The upper panel shows results for testing each feature individually, whereas the lower panel displays association statistics accounting for distance to gene footprint. **b** Predicted probability with 95%-confidence interval of distance to footprint across the five different quasi-true-positive sets. Estimates are from logistic regression models, modelling distance using restricted cubic splines. **c/d** Same as b, but modelling distance in the knowledge graph between gene and outcome (**c**) and CADD score (**d**). **e** Odds ratios from logistic regression models, ascertaining the association of the quasi-true-positive gene in the locus being a drug target.

We systematically tested for possible explanations of these differences based on gene and regional features, including putative effector genes being under differential selection pressure, which may extend to greater conservation of surrounding regulatory regions, and measures of regulatory complexity within the region (**Supplementary Tab. 5**). These features, however, explained the differences only partially. For example, while genes included in the mQTL-derived dataset showed the lowest evolutionary constraint, approximately 1.38 Z-score units lower (p<4.1×10^-24^) than putative effector genes included in the PoPS-derived dataset, other candidate gene sets showed minor differences (**Supplementary Fig. 3**). Similarly, only the number of recombination hotspots (>10 cM/Mb) between the lead variant and the putative effector gene differed significantly (p<7.5×10^-10^) across quasi-true-positive sets but differences were minor (e.g., mQTL effector genes had on average 0.49 fewer hotspots in between) and unlikely to account for the differential predictions.

Collectively, these results indicated a fundamental difference in the complexity of local genetic control of gene sets across quasi-true-positive sets that current feature sets do not fully capture.

### Divergent gene assignment methods recover known biology but poorly agree on novel insights

To evaluate the extent to which effector gene annotation strategies influence the biological interpretation of GWAS results, we next annotated lead variants for six common polygenic diseases: type 2 diabetes, ischaemic heart disease, hypothyroidism, prostate cancer, rheumatoid arthritis and schizophrenia^16–21^; **Tab. 1** and **Supplementary Tab. 6**). In line with benchmarking results, we observed considerable heterogeneity in effector gene assignments with only 17.1% (type 2 diabetes) to 31.8% (rheumatoid arthritis) of effector genes shared among all trained classifiers (Fig. 4a). Although agreement improved when omitting single classifiers, e.g., 31.6% - 45.9% when imposing agreement among four out of five classifiers, our results corroborate the need for standardised workflows and reporting of effector gene assignment^6^. We further report nearest-gene assignment by transcription start site or regulatory footprint in the following as a benchmark, since it encodes genomic proximity rather than learned functional evidence.

**Fig. 4.**
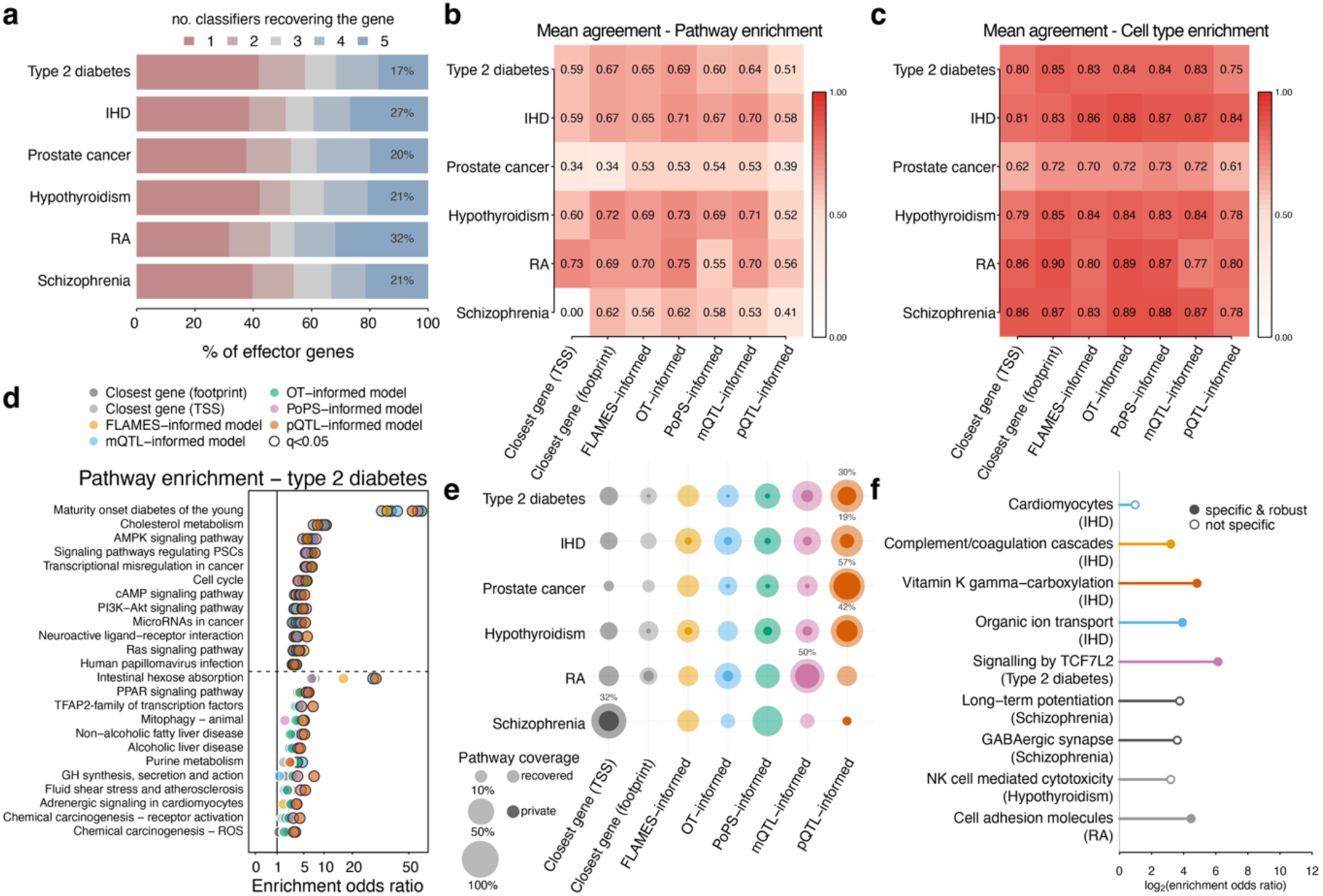
Heterogeneity in effector gene assignment propagates to the biological interpretation of GWAS. Lead variants from six European-ancestry GWAS of common polygenic disease (Tab. 1) were assigned effector genes by seven variant-to-gene classifiers: two genomic-proximity baselines (closest gene by transcription start site (TSS) or regulatory footprint) and five trained models informed by FLAMES, mQTL, Open Targets (OT), PoPS or trans-pQTL data. **a**, Sharedness of effector gene assignments across the five trained classifiers. Bars give the percentage of all genes assigned for a disease that were assigned by exactly one to five classifiers (rose to blue); the annotation gives the percentage assigned by all five. **b,c,** Mean pairwise agreement between classifiers for pathway (b) and cell-type (c) enrichment, computed as the mean off-diagonal Spearman rank correlation of a signed enrichment score (−log₁₀(FDR) × sign (odds ratio − 1)) across the union of tested terms; each cell gives one classifier’s mean agreement with the remaining six for that disease. The value of 0.00 for closest gene (TSS) in schizophrenia reflects the absence of any significantly enriched pathway for that classifier, not maximal disagreement. **d**, Within-disease coherence of pathway enrichment for type 2 diabetes. Rows are representative terms of redundancy-collapsed pathway modules (the most significantly enriched member of each module); one dot per classifier gives its multifunctionality-corrected enrichment odds ratio on a square-root axis, with dark outlines marking FDR < 0.05. Terms above the dashed line were enriched in at least five of seven classifiers (consistent), those below in two to four (divergent); up to 12 terms are shown per group, ranked by the number of classifiers and then by mean odds ratio. PPAR signalling, discussed in the text, is shown in addition and replaces the divergent term with the lowest mean odds ratio. **e**, Pathways enriched for a single classifier, shown relative to what each classifier recovered. Pale circles give the number of pathways significantly enriched for that classifier and disease as a percentage of all pathways enriched for any classifier in that disease; solid circles give the subset enriched for no other classifier. Area is proportional to percentage. Annotations give the share of a classifier’s enriched pathways that no other classifier recovered, for the most distinct classifier per disease, where that classifier enriched at least ten pathways. **f**, Pathways and cell types significantly enriched for exactly one classifier and discussed in the text, shown as log₂ enrichment odds ratio and coloured by that classifier as in d and e. Filled circles denote terms that were both specific and robust to multifunctionality correction, open circles those that were not. Throughout, enrichment was tested by one-sided Fisher’s exact test against KEGG and Reactome pathways or Human Protein Atlas v25 single-cell marker sets, with multifunctionality correction and Benjamini–Hochberg control; terms were considered enriched at FDR < 0.05 with an odds ratio > 1 and at least three overlapping genes. IHD, ischaemic heart disease; RA, rheumatoid arthritis

**Table 1.** GWAS of six distinct human phenotypes in populations of European ancestry.

| **Phenotype** | **Entry** | **N cases\*** | **Number of lead variants** |
|---|---|---|---|
| **Ischaemic heart disease (IHD)** | GCST90132314 | 181,522 | 317 |
| **Type 2 diabetes (T2D)** | GCST90492734 | 242,283 | 885 |
| **Schizophrenia** | GCST90128471 | 53,386 | 231 |
| **Hypothyroidism** | GCST90572791 | 98,805 | 177 |
| **Prostate Cancer** | GCST90274714 | 122,188 | 305 |
| **Rheumatoid arthritis** | GCST90132223 | 22,350 | 51 |

The observed heterogeneity at the gene level propagated to results from pathway (**Fig. 4b**) and to a lesser extent cell-type enrichment (**Fig. 4c**). All classifiers agreed only moderately on the enriched pathways (mean pairwise rank agreement ρ=0.59; range 0.46-0.67 across the six diseases) but more strongly at the level of cell types (ρ=0.81; range 0.69-0.85). The pQTL-informed model produced the most distinct results, consistent with its reliance on distal, protein-level regulation rather than genomic proximity or local functional genomics, and thus recovered partly non-overlapping effector genes and downstream enrichments. In detail, only 12.9% of enriched pathway modules were shared by all seven classifiers, against 29.5% of cell types, whereas 36.1% of pathway modules were enriched among gene sets from a single classifier, compared with 25.0% of cell types.

The observed heterogeneity mattered most where GWAS may yield actionable insights into disease processes. Core biological pathways and relevant cell types, by contrast, were recovered by most classifiers, albeit with notable exceptions and widely varying enrichment scores (**Supplementary Tabs. 7 and 8**). For example, PPAR signalling in type 2 diabetes was recovered by only four of the seven classifiers, the footprint- (OR=5.4; FDR=4.5×10^-2^), mQTL- (OR=5.8; FDR=1.2×10^-2^), PoPS- (OR=5.6; FDR=7.9×10^-3^) and pQTL-informed (OR=6.1; FDR=4.3×10^-3^) models, but did not reach significance with the FLAMES- or Open Targets-informed models (**Fig. 4d**). The implicated genes comprised canonical pathway members such as the nuclear receptors *PPARA*, *PPARG*, and their heterodimer partner *RXRA*, together with downstream targets integral to lipid handling, such as *ADIPOQ*, *ANGPTL4*, *LPL*, and *APOC3*. This pathway is of direct translational relevance: PPARγ, encoded by *PPARG*, is the target of thiazolidinediones and PPARα, encoded by *PPARA*, of fibrates^22^, and saturation mutagenesis of PPARG established that loss-of-function alleles cause Mendelian lipodystrophy and increase type 2 diabetes risk^23^.

Beyond graded disagreement, 185 pathways and 33 cell types were recovered by exactly one of the seven classifiers. Because classifiers differed in how many enriched terms reached significance, we expressed these private findings relative to each classifier’s own yield within a disease (**Fig. 4e**). This normalisation confirmed that exclusivity was a property of the classifier rather than a function of the number of significantly enriched pathways.

For example, 57% of the pathways significantly enriched among effector gene sets based on the pQTL-informed model in prostate cancer were private (47 of 82), and 42% in hypothyroidism (27 of 64), with similar patterns across the remaining diseases and classifiers (Fig. 4e), whereas the proximity baselines contributed few private pathways overall, with one striking exception discussed below. Critically, those findings included multiple examples of genuine disease biology that would not have been reported had the analysis relied on any other single classifier (**Fig. 4f**). In most instances, however, the remaining classifiers assigned the same driver genes with comparable enrichment that did not survive correction for multiple testing; only 28 of the 185 private pathways, 27 of them from the pQTL-informed model, showed no nominal enrichment (P≥0.05) in any other classifier. Heterogeneity therefore reflected scattered statistical support more than divergent gene content, motivating aggregation of evidence across classifiers (see next section).

Genes with enhanced expression in cardiomyocytes were uniquely enriched among effector genes for ischaemic heart disease in the mQTL-informed model (OR=1.97; FDR=4.5×10-2), driven by canonical coronary risk genes alongside genes central to cardiomyocyte identity and cardiac development^24^ (Supplementary Note 1). The complement and coagulation cascade as a pathway possibly predisposing to ischaemic heart disease was likewise enriched only using results from the FLAMES-informed classifier (OR=9.2; FDR=2.4×10^-2^). The enrichment reflected canonical constituents of both arms: the classical-pathway complement protease C1S together with the coagulation and fibrinolytic regulators *F10*, *PLG*, *PROCR*, and *SERPINA1*. Pharmacological suppression of coagulation downstream of F10 has proven interventional benefit, reducing major ischaemic events by about 24% in the COMPASS trial^25^. Evidence for the complement arm, in contrast, remains preclinical, spanning complement deposition in unstable human atherosclerotic lesions^26^, attenuated atherosclerosis and ischaemia– reperfusion injury upon complement inhibition in experimental models^27–29^, and C1-inhibitor as a proposed therapeutic candidate without definitive trial evidence^30^. Two further private examples for ischaemic heart disease, vitamin K-dependent γ-carboxylation and organic ion transport, are detailed in Supplementary Note 1.

For type 2 diabetes, the strongest private signal was Wnt signalling through *TCF7L2* (*TCF7L2*, *TCF4*, *CTBP1*; PoPS-informed model; OR=70.6; FDR=4.7×10^-3^), despite TCF7L2 harbouring the strongest common risk variant for the disease. Illustrating the point above, five of the six other classifiers assigned the identical three genes with comparable enrichment (OR=13.3-34.9), but none passed correction.

Even trained on rich functional genomic data, V2G models did not necessarily report coherent disease biology. Nearest-gene assignment by regulatory footprint showed the highest fraction of uniquely enriched pathways for schizophrenia (6 of 19 pathways, 32%), and those implicated well established disease biology. For example, long-term potentiation that strengthens synaptic ties among neurons (*GRIN2A*, *GRIA1*, *ERBB4*, and *NRGN*; OR=13.5; FDR=4.3×10^-2^), and GABAergic synapse signalling, calcium signalling and dopaminergic transmission (*DRD2*, *GRIN2A*, and *GRM3*) would not have been reported from any trained classifier, despite GRIN2A carrying convergent common- and rare-variant evidence^21,31^ (Supplementary Note 1). All five trained classifiers nonetheless assigned at least three of the four driver genes of long-term potentiation, with comparable enrichment (OR=10.3-13.2; nominal P≤3.8×10^-3^) that did not survive multiple testing correction (FDR≥6.4×10^-2^). Across the remaining diseases, the two proximity classifiers contributed three private pathways in total, two of them with genuine disease relevance: natural killer cell mediated cytotoxicity in hypothyroidism (*CD247, IFNG,* and *ZAP70*; OR=9.2; FDR=3.7×10^-2^) and cell adhesion molecules in rheumatoid arthritis (*CD2, CD40, CTLA4,* and *ICAM3*; OR=22.1; FDR=4.8×10⁻²), the latter recovered only by nearest-gene assignment by transcription start site. *CTLA4* encodes the target of the costimulation modulator abatacept, which improved clinical response in rheumatoid arthritis refractory to TNF inhibition^32^ (Supplementary Note 1).

Collectively, these results illustrated that coherent recovery of known biological pathways and cell types is insufficient to benchmark effector gene assignment methods from which novel biology is subsequently inferred. For example, an analysis relying on a classifier selected for its aggregated performance would not have reported either the complement and coagulation cascade in ischaemic heart disease or long-term potentiation in schizophrenia, and no single classifier reported the full landscape of disease relevant biology corroborated by GWAS.

### Successful recovery of established drug targets is classifier-dependent

To evaluate the impact of different effector gene prioritisation frameworks on translational utility, we assessed their capacity to recover established drug targets supported by human genetic evidence, used here as an orthogonal evaluation set rather than as ground truth. We assembled 340 locus-target-indication triplets in which an approved drug target lay within a 1-Mb window of the lead variant for the indication and scored every candidate gene at each locus (**Supplementary Tab. 9**). Recovery was strongly classifier-dependent (Cochran’s Q = 88.4, d.f. = 6, P = 6.4×10^-17^) and classifiers differed in average performance (Friedman P = 3.2×10^-38^; **Fig. 5a**). The PoPS-informed model achieved the highest mean within-locus discrimination (AUC = 0.80; target ranked first at 38.8% and within the top three at 62.4% of loci), followed by the Open Targets-(within-locus AUC = 0.76) and pQTL-informed (within-locus AUC = 0.75) models, whereas nearest-gene assignment reached an AUC of only 0.64. Consistent with the gene-level heterogeneity observed in enrichment analysis, the pQTL-informed model ranked the target first most frequently of all classifiers (41.5% of loci) despite its lower overall AUC, indicating that it identified the target gene confidently at some loci while missing it at others.

**Fig. 5.**
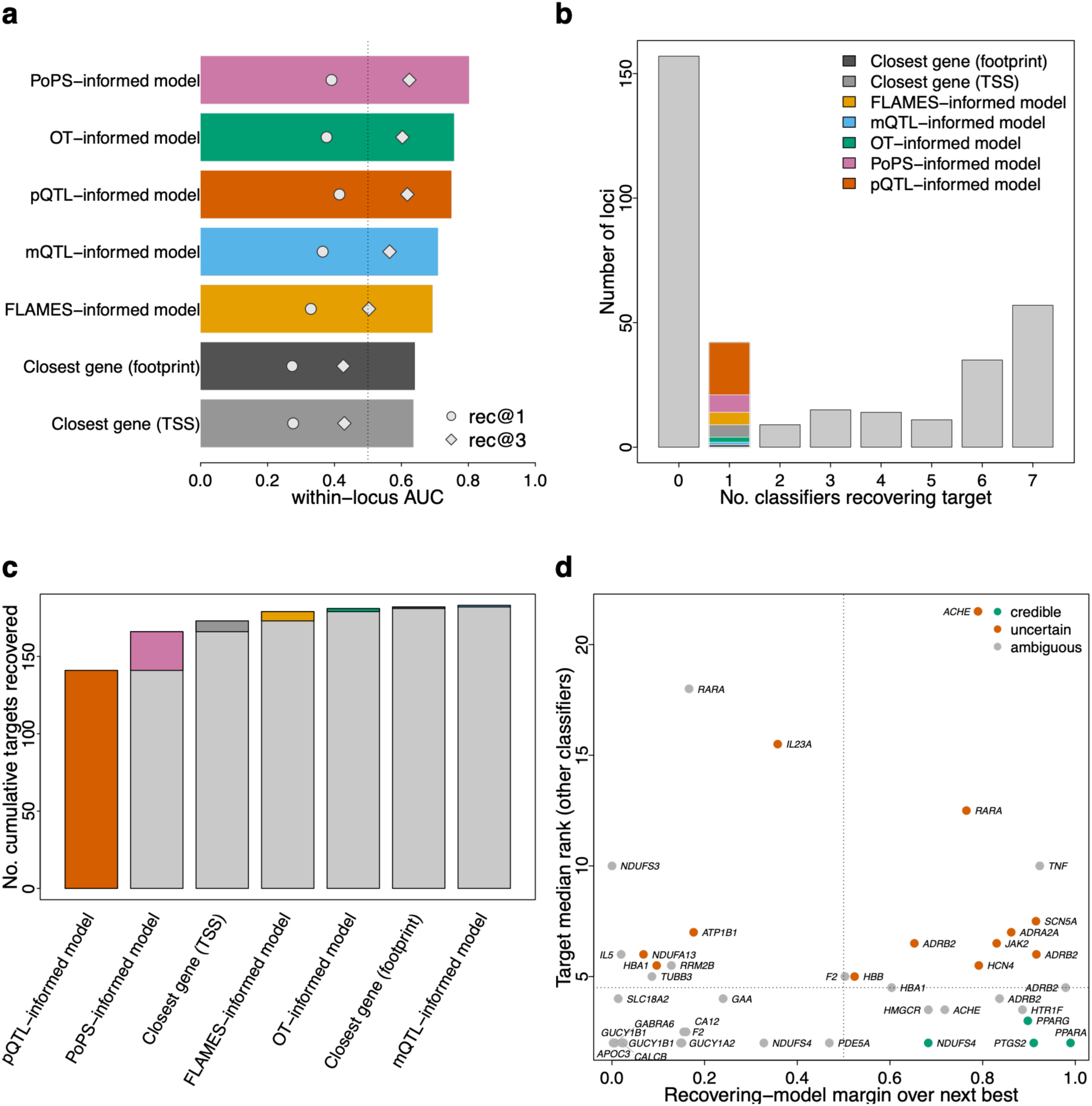
Recovery of established drug targets by variant-to-gene classifiers is heterogeneous and classifier-dependent. Recovery of the validated drug target among all protein-coding candidate genes within 1 Mb of the lead variant, across 340 locus–target–indication triplets. **a,** Per-classifier recovery performance, ordered by within-locus AUC (bars; the probability that the target outranks a randomly chosen competing gene at the same locus). Points show the proportion of loci at which the target was ranked first (rec@1) or within the top three (rec@3). b, Distribution of the number of classifiers recovering each target as the top-ranked gene (0–7) **c,** Greedy cumulative recovery, in which classifiers are added in order of their marginal gain in newly recovered targets. **d,** Audit of the 42 targets recovered by a single classifier, showing the recovering model’s score margin over the next-best candidate at the locus (x axis) against the median rank assigned to the target by the remaining six classifiers (y axis). Points were coloured by credibility (credible, uncertain, ambiguous) and labelled by target gene. Targets combining a large recovering-model margin with a high median rank elsewhere (upper right; for example, SCN5A and ADRB2) are prioritised by a single model against the consensus of the others, whereas credible single-model recoveries (for example, PPARA, PPARG, PTGS2 and NDUFS4) retain a low median rank across classifiers.

We observed high heterogeneity in scoring behaviour across the loci. About half (n=188) of the 340 targets were recovered as the top-ranked candidate by at least one of the seven classifiers (n=177 by the five domain-informed models alone), but only 57 (16.8%) were recovered unanimously by all seven classifiers (**Fig. 5b**). Agreement among the seven classifiers’ per-locus rankings was low (Kendall’s *W* = 0.064, P = 8.4×10^-26^), indicating that the best classifier for identifying the causal gene differed markedly from locus to locus rather than reflecting a single, uniformly superior approach. The pQTL-informed model recovered most loci (n=141), with the PoPS-informed model capturing a further 25 (cumulative 166, 91% of the union), while the remaining classifiers each contributed at most seven additional targets (**Fig. 5c**). Notably, nearest-gene assignment by transcription start site recovered five targets missed by all five trained models.

Since a target recovered by a single classifier may reflect either a uniquely captured signal or a misassignment, we audited the 42 loci recovered by exactly one classifier (**Fig. 5d**). The pQTL-informed model was responsible for half of all single-model recoveries (21 of 42), including eleven of uncertain significance, such as *ADRB2*, *ADRA2A*, *SCN5A*, *HCN4*, *JAK2*, *ACHE*, and *RARA*, that were ranked low by all other classifiers and showed only minimal higher predicted probabilities than the next best gene in the locus, possibly reflecting the tendency of the classifier to rank well-curated protein interaction partners highly irrespective of distance to the lead variant at the locus. Conversely, credible single-model recoveries included *PPARA*, *PPARG*, and *PTGS2* by the pQTL-informed and *NDUFS4* by the PoPS-informed model, representing genuine rescues of the correct gene that the other classifiers underweighted. The recovery of *PPARA*, encoding the target of fibrates, and *PPARG*, encoding the target of thiazolidinediones, is noteworthy, as both encode ligand-activated nuclear receptors with extensive experimental evidence for their roles in lipid metabolism and lipoprotein regulation (PPARα; fibrates for lipid disorders) and in adipocyte lipid storage and insulin sensitisation improving glucose homeostasis^22^ (PPARγ; thiazolidinediones for type 2 diabetes).

Collectively, these results demonstrate that recovery of established drug targets was strongly classifier-dependent, and that single-model nominations require corroboration before being treated as high-confidence effector genes.

### Consensus models matched domain-specialist performance at the cost of uniquely recovered drug targets

Having shown that domain-specific models inferred partially distinct biology and recovered different druggable targets, we asked whether combining domains within a single training dataset would be able to consolidate across those differences. We therefore pooled the training data from the distinct models into one consensus model. On average, this model reached in-domain performance statistically indistinguishable from domain-specific models (**Fig. 6a**). Its recovery of drug targets, however, was worse (124 out of 340, 36.5%) than almost every individual classifier, surpassing only the FLAMES-informed model (**Fig. 6b**), demonstrating that simply pooling or diversifying training data is not sufficient to achieve equitable performance.

**Fig. 6.**
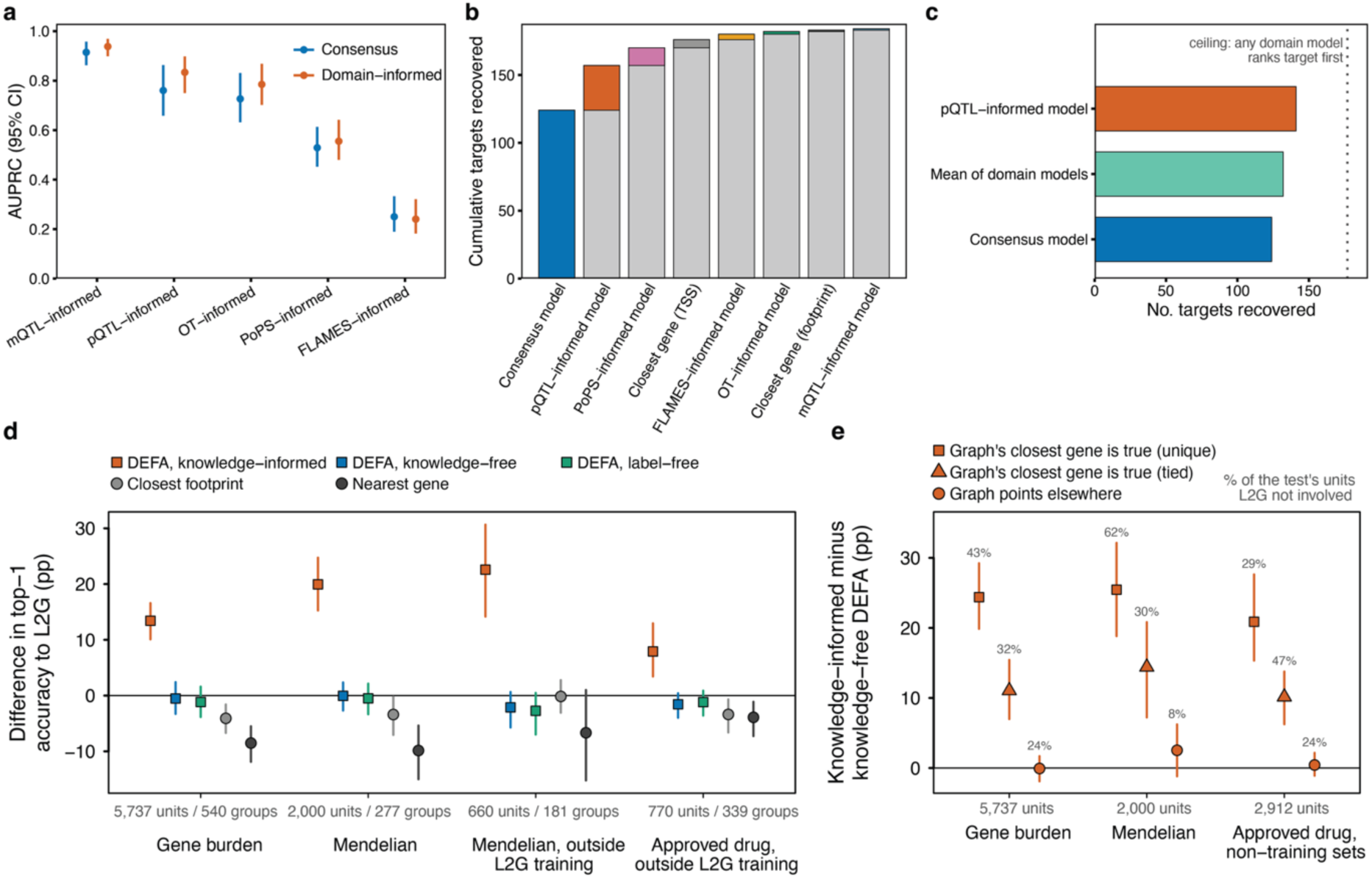
Combining domain-informed models outperforms pooled training for drug-target recovery, and DEFA matches or exceeds Open Targets L2G. a, In-domain performance (AUPRC, 95% CI) of a consensus model trained on the pooled five quasi-true-positive sets, compared with the domain-informed model trained on each set alone. b, Greedy cumulative recovery of the 340 approved drug targets, starting from the consensus model and adding classifiers in order of their marginal gain; coloured segments mark targets newly recovered by each classifier. c, Strategies for combining models: averaging the scores of the five domain-informed models recovers more targets as the top-ranked gene (n=132) than the consensus model (n=124), but not more than the best single model, the pQTL-informed model (n=141). The dotted line marks the ceiling of targets ranked first by at least one domain-informed model (n=177). d,e, DEFA compared with Open Targets L2G on credible set–trait pairs scored by both methods, against gene-burden associations (5,737 pairs in 540 gene–disease groups), Mendelian gene–disease relations (2,000 pairs, 277 groups; and 660 pairs, 181 groups, outside L2G training) and approved drugs for the same disease outside L2G training (770 pairs, 339 groups). DEFA assigned the gene with the highest mean probability across its classifiers. Pairs outside L2G training are those whose credible set is not an L2G training positive and whose locus and gene–disease relation do not overlap one. d, Difference in top-1 accuracy to L2G (percentage points; 95% CI from a paired bootstrap over linked gene–disease groups, 2,000 resamples) for knowledge-informed DEFA, knowledge-free DEFA, its label-free version without the Open Targets-informed model, the closest gene by regulatory footprint and the nearest gene. e, Accuracy gain of knowledge-informed over knowledge-free DEFA, stratified by whether the true gene was the only candidate closest to the trait in the knowledge graph, tied with other candidates, or not the closest; points give the gain (95% CI, same bootstrap) and labels the share of pairs in each stratum. This contrast does not involve L2G, so the approved-drug test uses all 2,912 pairs not on L2G training credible sets. AUPRC, area under the precision– recall curve.

We therefore next asked which strategy for combining probabilities across classifiers maximised drug-target recovery to guide consolidation for future efforts. Taking the mean relative score across the five domain-informed classifiers recovered 38.8% (n=132) of established targets, outperforming the consensus model (36.5%; n=124) but not the best individual classifier, the pQTL-informed model (41.5%; n=141) (**Fig. 6c**). All strategies fell short of the 177 targets (52.1%) ranked first by at least one domain-informed model, indicating that each curation assumption contributed targets that were lost on consolidation (**Fig. 6c**). Together, these results showed that no way of combining the five domain-informed models recovered what they nominated collectively. We nevertheless suggest averaging their scores over training a single model on all curations, because it recovered more targets and kept each nomination visible, so that agreement between models indicates confidence and disagreement points back to the curation assumption behind a prediction.

### DEFA matches L2G without external knowledge and gains where knowledge links a candidate gene to the trait

To situate our framework within the field, we finally benchmarked DEFA against Open Targets locus-to-gene^13^ (L2G) on 178,518 credible set–trait pairs scored by both methods, using three lines of evidence independent of the locus statistics used by either: rare-variant gene-burden associations (5,737 pairs in 540 independent gene-disease groups), Mendelian gene-disease relations (2,000 pairs; 277 groups) and approved drugs for the same disease at pairs outside L2G training (770 pairs; 339 groups; Methods). Without knowledge-graph features, DEFA was as accurate as L2G on all three tests (56.1% vs 56.7%, 58.5% vs 58.5% and 24.7% vs 26.2%; differences -1.6 to +0.0 percentage points, all 95% CIs including zero; paired bootstrap over gene–disease groups), exceeded nearest-gene assignment by 8-10 points on gene-burden and Mendelian genes, and did so even without the Open Targets-informed model. With knowledge-graph features, DEFA exceeded L2G by 13.4 (95% CI 10.1-16.6), 20.0 (15.3-24.7) and 7.9 (3.4-13.0) points, respectively, irrespective of how classifiers were aggregated (**Fig. 6d; Supplementary Tab. 10**).

We incorporated graph features to extend the biologically informed prior that guides annotations for molecular QTLs to complex diseases, and the performance gain was hence confined to pairs where the knowledge graph already linked a candidate gene to the trait (**Fig. 6e; Supplementary Tab. 11**). Where the true gene was the only candidate closest to the trait in the graph (29-62% of pairs across tests), knowledge-informed DEFA exceeded its knowledge-free counterpart by 20.9-25.5 points, and where it tied with other candidates (30-47%), by 10.2–14.4 points. Where the graph pointed to entirely different genes (8-24% of pairs), the gain was absent (all 95% CIs including zero), while accuracy remained on par with L2G (-4.4 to +4.2). Because Mendelian genes and approved-drug targets are themselves curated relations the graph encodes, graph distance alone outperformed the full model on these two tests, whereas on gene-burden genes, which overlap least with curated knowledge, combining prior and locus evidence (70.1%) outperformed both the prior alone (62.3%) and the knowledge-free model (56.1%; **Supplementary Fig. 4**). For genes first reported after the graph’s mid-2021 sources, too few independent gene-disease groups remained to test whether this lead persisted (Mendelian genes: +5.8 points, 95% CI -3.6 to 19.5, 172 pairs in 28 groups; approved-drug targets: +7.8, -25.5 to 27.9, 153 pairs in 23 groups; **Supplementary Tab. 11**).

## Discussion

Effector gene assignment remains the principal bottleneck in converting (non-coding) GWAS signals into testable biology. Here we showed that 1) the poor transferability of V2G models across biological trait domains reflected domain-specific regulatory architecture rather than model architecture, with genomic distance carrying the most divergent weight of 269 features tested (p_interaction_<5.3×10^-123^; **Fig. 3b**); 2) the biological assumptions embedded in curating quasi-true-positive sets propagated into predictions and can become self-confirming; 3) aggregating evidence across domain-specific classifiers, rather than pooling training data, provided superior recovery of established drug targets (n=132 versus n=124); and 4) DEFA matched Open Targets L2G without external knowledge and exceeded it where curated biology links a candidate gene to the trait (Fig. 7).

GWAS remain the least biased readout of causal biology in living humans, capturing millions of natural experiments that link perturbed gene function to disease in the general population. Single genetic variants, however, explain very little variation in complex phenotypes^33^, and inference on the role of pathways, cell types, mechanisms and therapeutic hypotheses rests on the identification of the correct effector gene. Models predicting effector genes have, in turn, been trained on curated knowledge that is both incomplete and unevenly distributed across biology. Classifiers accordingly agreed on established disease biology, yet shared only 17.1% to 31.8% of effector gene assignments across six common diseases, diverging precisely where GWAS are most informative. Our contribution is therefore not merely a better classifier, but a demonstration that the choice of assumption is consequential, quantifiable, and currently unreported.

Poor generalisability of models across independently ascertained domains may appear unsurprising. Our systematic interaction testing across more than 250 functional genomic features, however, suggested a more specific explanation: differential weighting of genomic distance, rather than model architecture, was the dominant driver. This pointed towards regulatory contexts and mechanisms that differ across curated biological domains. Adjusting for the differential association of distance strongly attenuated interaction statistics for most other features (**Fig. 3a**), consistent with distance acting as the primary axis along which domains separated. This interpretation was supported by independent observations from other modalities. Cell-type-resolved eQTLs are less likely to regulate the nearest gene than their tissue-level counterparts^34^, while enhancer - gene encyclopaedias report regulatory complexity varying substantially between genes^35^. We therefore attribute the poor transferability to differences in regulatory biology that current feature sets capture only incompletely, rather than to a failure of any individual method.

A second, more consequential source of divergence lay beyond genuine biological heterogeneity. The assumptions used to curate training data propagated into the predictions of the resulting models. The clearest example concerned druggability, which was the strongest single feature in the Open Targets-informed model (OR: 1.93, p<3.2×10^-98^) and effectively uninformative in the FLAMES-informed model (OR: 1.05, p=0.22). Because the Open Targets gold standard was curated in part from drug-target and rare-disease evidence^13^, a model trained on it learned features that identify drug targets and will subsequently appear to corroborate those, a loop that might be misread as biological validation. An analogous pattern applied to training sets guided by exonic variants^8,9^: the coding variant unambiguously identifies a gene, but the neighbouring non-coding signal need not act through that same gene, as mechanisms acting through non-coding variation have been shown to systematically prioritise different genes^36,37^. Coding variants thus offer an attractive anchor for curation precisely because they are unambiguous, but that unambiguity applies to the variant, not to the regulatory relationship it is used to represent. Each curation choice was well justified in the absence of ground truth, but their consequences remained invisible to benchmarks that reward recovery of established biology. Making the underlying assumption explicit, and evaluating models against domains built on deliberately independent reasoning, is in our view the most immediately actionable conceptual shift arising from our work.

Incorporating prior biological knowledge was essential to constructing diverse quasi-true-positive sets, and we further found that integrating such knowledge directly into classifiers improved cross-validated performance by 16.5% on average. Conceptually, this placed our approach close to top-down methods for gene prioritisation such as PoPS^8^, in which genome-wide polygenic signal informs annotation at individual loci, while making more explicit use of disease-related biology. However, even with extensive molecular evidence and a knowledge prior, many loci remained unresolved, including some whose biology is partly understood. Where the knowledge graph pointed away from the true gene, no method resolved these loci. Every method, including L2G, identified the approved-drug target in fewer than one in ten pairs (5.5-6.5%) and the gene-burden gene in fewer than half (37.7-44.0%). Even a tie between the true gene and other candidates left knowledge-informed DEFA at only 52-64% accuracy. At such loci prior knowledge narrowed the candidates without providing strong evidence for one over the others, and neither functional genomic annotation nor knowledge-graph evidence currently resolves which gene mediates the association.

Our results argued against committing to a single assumption before annotating effector genes, but they also indicated that pooling diluted rather than integrated domain-specific evidence. Taking the maximum score across the five domain-informed classifiers maximised drug target recovery, but strategies favouring sensitivity are prone to false positives that require independent corroboration, which cannot currently be automated. Scoring loci with several models makes disagreement itself informative, since a confident assignment and a confident error are otherwise hard to distinguish. We therefore consider DEFA, which we make freely available (https://github.com/lkohleick/DEFA), a partial and pragmatic answer to current calls for standardised V2G workflows and reporting^6^.

Several limitations need to be considered when interpreting our conclusions. First, our three evaluation sets were themselves quasi-true positives, modest in size (n=62, 29 and 101), and rested on curation rather than experiment; their use for a transferability test rested on the independence of those lines of evidence, not the correctness of any one. Second, curated datasets imposed a single effector gene per locus based on established biology, a framework challenged by emerging evidence that individual regulatory elements act on several nearby genes^35^, and predictions returning calibrated sets of candidate genes rather than a single winner would be a natural extension. Third, our candidate space comprised protein-coding genes within a fixed window, so loci acting through long non-coding RNAs or other regulatory moieties are missing. Fourth, the knowledge graph we used encoded curated disease-gene relationships, which may have inflated performance for well-studied genes; we therefore reported results with and without graph-derived features. Fifth, our domains were defined through molecular evidence (mQTLs and trans-pQTLs), an approach that is tractable for such but does not readily extend to diseases, for which the relevant regulatory mechanisms remain often unknown. Hence, delineating disease-relevant domains is in our view the principal open problem arising from our work. Sixth, recovery of approved drug targets, while translationally meaningful, is a biased benchmark, and future models will need training against experimental evidence. Finally, all GWAS analysed were of European ancestry, and because distance was our dominant differential feature, differences in linkage disequilibrium across ancestries are likely to exacerbate misalignment.

Most of the named limitations are due to the absence of an unselected, experimentally resolved set of variant-gene pairs spanning the breadth of regulatory mechanisms in the human genome. Until such a reference exists, every V2G model, including ours, encodes an assumption about how variants act on genes. Future models could treat the closest gene as an explicit null hypothesis and quantify how strongly the evidence at a locus supports a more distal candidate. The preference for nearby genes, currently built into every method without being stated, would then become a testable assumption, while a single generalist model could still adapt to distinct locus subtypes. Improved representation learning, including interpretable path-based architectures for knowledge graphs^38^, may strengthen the augmentation reported here while making its contribution explainable. Sequence-based models of regulatory variant effect^39^ and cell-type-resolved QTL mapping offer corroboration independent of curated gene sets, though increased colocalisation reflects regulatory resolution rather than causal involvement of the gene in disease onset^34^.

## ACKNOWLEDGEMENTS

The work was funded by the European Union (ERC, GenDrug, 101116072). Views and opinions expressed are however those of the author(s) only and do not necessarily reflect those of the European Union or the European Research Council. Neither the European Union nor the granting authority can be held responsible for them. The funders had no role in study design, data collection and analysis, decision to publish or preparation of the manuscript. We thank Karl Heilbron for discussion and valuable input. We acknowledge the Scientific Computing of the IT Division at the Charité – Universitätsmedizin Berlin for providing computational resources that have contributed to the research results reported in this article. We used LLM-based tools for software development and analyses, with close supervision and manual review. This research has been conducted using the UK Biobank Resource under Application Number 44448

## DATA AVAILABILITY

Associated data will be made available upon final publication.

## CODE AVAILABILITY

DEFA is available via GitHub (https://github.com/lkohleick/DEFA). Code covering the analysis will be deposited on GitHub upon final publication.

## AUTHOR CONTRIBUTIONS

Conceptualization: LK, BW, MP

Data curation/Software: LK, KH, CB, MZ, MP

Formal Analysis: LK, MP

Methodology: LK, BW, MP Visualization: LK, MP

Funding acquisition: CL, MP

Project administration: MP

Supervision: MP

Writing - original draft: LK, MP

Writing - review & editing: all

## Methods

### Construction of high-confidence evaluation datasets

We first sought to leverage the architecture of protein quantitative trait loci (pQTLs) to disentangle genetic effects on circulating protein levels. By identifying variants associated with protein levels in *trans*, we hypothesised that local genetic effects at the variant’s locus were unlikely to be secondary to the modulated *trans*-gene (Fig 1c) but rather were affecting an interaction partner of the protein in the locus. Applying this logic to ligand-receptor pairs from OmniPathR^40^ and protein complex data from CORUM^41^ and huMap 2^42^, and after removing pleiotropic loci, removing loci in intronic regions and further extensive manual curation, we identified 101 high-confidence fine-mapped credible sets with assigned effector genes. Information on trans-pQTLs and corresponding summary statistics were obtained from the UKBB Pharmacoproteomics resource^43^.

In a parallel effort, we integrated findings from recent large-scale metabolomics studies to define a metabolite-based allelic series. We selected credible sets where gene assignments in the UK Biobank (UKBB) trans-ancestral map matched those reported in parallel independent efforts for colocalizing loci^44,45^. To mitigate concerns regarding nonspecific pleiotropy, we excluded all overlapping credible sets of lipid-related metabolites, resulting in 62 high-confidence mQTL-to-gene assignments.

Finally, we harnessed the link between human disease genetics and drug-target biology to generate 29 credible sets where the GWAS signal for a specific indication matched an approved drug target within 750 kb (Fig. 1b) with available summary statistics and extensive manual review.

### Benchmarking Metrics

Performance was assessed using the area under the precision-recall curve (AUPRC) to account for the class imbalance inherent in variant-to-gene prioritisation (one effector gene vs. many neighbouring genes). Additionally, to estimate recovery of established drug targets, we used recall at k=1 and k=3 (rec@1 and rec@3), which measures the recall of relevant items in a dataset in the top-k recommendations.

### Creating five distinct variant-to-gene datasets for model training

We assembled five complementary locus-gene datasets spanning proteomic, metabolite, GWAS-informed and curated gene-prioritisation resources. For training, we chose a deliberately smaller window of ±500kb than theoretically predicted^46^ to avoid enrichment of false-positive assignments during training and model evaluation.

#### pQTL-informed dataset

Similar to the benchmarking set, we set out to create a training dataset based on trans-pQTL signals. Relaxing the pleiotropy filters and less stringent manual curation yielded 416 loci with an assigned effector gene. Notably, effector genes were inferred only for this trans-pQTL dataset; in all other datasets, effector gene assignments were taken directly from the source resource.

#### mQTL-informed dataset

We obtained variant-gene assignments for metabolite QTLs from a study employing broad capture mass spectrometry^47^ resulting in 489 loci with an assigned effector gene.

#### OT-informed dataset

We used the variant-gene dataset from the initial Open Targets Locus-to-Gene (L2G) model^13^, yielding 441 loci with an assigned effector gene.

#### FLAMES-informed dataset

We used the loci provided in the FLAMES data driven training dataset and selected the variant with the highest PIP per fine-mapped locus as the lead variant, resulting in 698 lead variants with an assigned effector gene. In this dataset, the ‘positive’ gene at each locus was the gene harbouring an independent ExWAS signal within the locus window.

#### PoPS-informed dataset

We used the PoPS evaluation dataset, which defined evaluation loci as non-coding fine-mapped credible sets that are non-overlapping with a nearby protein-coding fine-mapped signal for the same trait. In this dataset, the ‘positive’ gene at each locus was the gene harbouring the independent coding fine-mapped variant (PIP > 0.5). For model input, we represented each locus by selecting the variant with the highest PIP in the non-coding credible set, yielding 882 loci for training.

We defined the variant set at each locus by LD augmentation (r^2^>0.5) based on European ancestry-matched LD reference panel from UK Biobank of the lead variant rather than by statistical fine-mapping, for three reasons. First, credible sets are available for only a minority of published GWAS and require in-sample or well-matched LD, whereas LD augmentation is uniformly applicable across the heterogeneous studies underlying our evaluation and training sets. Second, fine-mapping from summary data is sensitive to inconsistency between association statistics and reference LD^48^, and calibration degrades further under the between-cohort heterogeneity intrinsic to meta-analysis. This has been shown across 14 large meta-analyses, in which 67% of loci showed patterns inconsistent with reliable fine-mapping, with suspicious loci depleted for nonsynonymous lead variants^49^. Annotation-derived features computed over credible sets would have therefore inherited an error structure that is itself correlated with functional annotation. Third, and most importantly for our design, credible-set width scales with statistical power, which differs by orders of magnitude across our biological domains. This may have otherwise introduced a power-dependent artefact for any distance-related investigations confounded by domain membership. LD augmentation, being a property of the reference panel rather than of study design, is invariant to power. We note that this choice trades precision for robustness, as the augmented set contains non-causal variants by construction, and we therefore report sensitivity to the r^2^ threshold in **Supplementary Fig. 6**. The domain-model mean is calculated by min-max scaling each of the five domain-informed models’ scores within the locus and then averaging these scaled scores for each candidate gene.

The UKBB was approved by the National Research Ethics Service Committee North West Multi-Centre Haydock and all study procedures were performed in accordance with the World Medical Association Declaration of Helsinki ethical principles for medical research. All analyses were conducted under UKBB application 44448.

### Annotation of genetic variants

We developed a modular variant annotation framework to characterise sentinel variants and their respective genomic loci. For each locus, the module incorporates either high-confidence proxies (defined by a user-specified r^2^ threshold) or variants contained within fine-mapped credible sets. Scores are calculated by multiplying the r^2^ for SNP-related annotations where applicable. This weighting mechanism generates a locus-specific annotation matrix, typically restricted to a 500kb window flanking the sentinel variant or the locus centre. All variants were harmonised internally to both GRCh37 and GRCh38 coordinates to enable consistent cross-resource integration.

Annotation features included VEP scores^33^ to capture the predicted functional consequence of each variant, and CADD scores to estimate variant deleteriousness^50^. To incorporate expression quantitative trait locus (eQTL) evidence at tissue-level resolution, we included fine-mapped eQTLs from the eQTL Catalogue Release 7, retaining the per-tissue signal^12^. To capture cell-type-resolved chromatin accessibility, we intersected variants with candidate cis-regulatory elements and their associated Activity-by-Contact (ABC) scores from the single-cell chromatin accessibility atlas (CATlas)^51^,

We further annotated each locus using the variant-to-gene annotation resource bundled with FLAMES^9^; the underlying data sources and their original references are listed in [Tab 1 of ref. FLAMES^9^]. These annotations comprised fine-mapped molecular QTL colocalisation evidence, an independent set of ABC enhancer–gene predictions together with a subset restricted to CRISPRi-validated enhancer–gene pairs, enhancer activity and enhancer–gene correlations derived from CAGE, PRO-seq/GRO-seq and integrative epigenomic data, physical chromatin evidence from promoter capture Hi-C and single-cell ATAC-seq co-accessibility, and promoter overlap. For each FLAMES-bundled resource, we intersected the variants at a locus with the annotated elements of every gene in the locus, weighted each overlap by the corresponding variant weight, and aggregated the resulting scores per locus–gene pair as their sum, maximum and, where applicable, mean.

Finally, we queried a list of approved drug targets from the Open Targets Platform^14^ (downloaded 2 July 2024; excluded during training of our domain-informed effector gene prediction models) and genes implicated in Mendelian disorders from OMIM^52^ (downloaded 10 April 2024).

#### Knowledge graph integration

To incorporate systemic biological context, we integrated graph-based features using the Precision Medicine Knowledge Graph (PrimeKG)^15^. When the GWAS endpoint was provided, we mapped the disease or molecular phenotype to its corresponding node within the graph to calculate connectivity metrics between the endpoint and candidate genes at each locus. Using the NetworkX package (version 3.5), we computed three primary topological features: the shortest path distance, to quantify the proximity of candidate genes to the disease node Personalised PageRank (PPR), to measure each gene’s relevance to the disease endpoint across the wider network; and the Adamic-Adar index, to identify genes sharing neighbours with the endpoint, weighting sparsely connected neighbours more heavily. We additionally generated these scores while excluding Gene Ontology (GO) features from the graph structure. The resulting annotation matrix provides a comprehensive multidimensional feature set for downstream effector gene prioritisation. A full list of annotation metrics is provided in **Supplementary Tab. 2**.

### Training of a flexible framework for effector gene assignment

We employed an extreme gradient boosting (XGBoost) framework to prioritise candidate effector genes based on the result of our annotation pipeline for each domain-informed dataset. To ensure robust generalisation and prevent over-fitting to specific genomic regions, we implemented a leave one-chromosome-out (LOCO) training procedure. For each dataset, one classifier was trained per autosome (chr1–chr22; chromosome X was additionally held out as a separate fold for the dataset in which it was present), each time training on the remaining chromosomes and evaluating on the held-out one. Hyperparameter optimisation was performed using a ten-iteration, five-fold cross-validation randomised grid search. One main XGBoost model per dataset was then trained using the mean hyperparameters of all LOCO models, refit on a random (8/9) of loci with the remaining (1/9) held out for early stopping and evaluation.

To evaluate generalisation across annotation sources, we additionally trained a pooled consensus model combining all five datasets. For each dataset, we held out a fixed 20% of loci (by variant identifier) and trained the LOCO procedure described above on the remaining 80%. The five 80% training splits were then pooled (removing duplicate variant–gene pairs) to train a single cross-dataset consensus model, using the same LOCO and hyperparameter-tuning procedure. Performance was evaluated within-dataset on each dataset’s own 20% holdout, and across-dataset by applying each dataset-specific model to the other four datasets in full.

As the number of loci was limited for several datasets, we estimated ranges of model performance (AUPRC) using a grouped bootstrap with replacement at the locus level (2,000 resamples, 95% percentile confidence intervals). This pipeline was run twice, with and without graph-derived (network-propagation) features, and a paired locus-level bootstrap was used to test whether the AUPRC difference attributable to graph features was distinguishable from zero.

### Model validation across different datasets

To evaluate the transferability of our approach, the pQTL-informed model was independently validated using a dataset of 91 pQTLs derived from the Fenland study using the SomaScan v4 assay^53^, reaching a performance of 0.88 AUPRC.

### Differential feature association testing

To understand poor transferability of models across quasi true-positive sets, we performed logistic regression analysis to test for a potential domain-by-feature interaction effect on odds ratios for the likely true positive gene in each locus. We used a flexible framework that allowed feature associations to be modelled non-linearly using restricted cubic splines with three knots. Splines were only used if likelihood ratio tests indicated a significantly better fit. We first tested each feature also included in machine learning models separately (univariate models) and subsequently repeated association testing accounting for non-linear interaction effect of distance to gene footprint. We considered interactions significant at p<1.8×10^-4^, a Bonferroni correction for the 269 features tested (**Supplementary Tab. 4**).

We further collated features describing evolutionary constraint of effector genes and complexity of the genomic region surrounding them^54^ (**Supplementary Tab. 5**). To characterise the regulatory architecture of each variant–gene pair, genomic features were derived from four complementary data sources. Candidate cis-regulatory element (cCRE) annotations, including promoter-like signatures (PLS), proximal and distal enhancer-like signatures (pELS/dELS), and CTCF-bound elements, were obtained from the ENCODE Registry of cCREs v4 (SCREEN), which catalogues 926,535 high-confidence elements across the human genome based on integrated DNase-seq, H3K4me3, H3K27ac, and CTCF ChIP-seq data from hundreds of biosamples^55^. For each variant-gene pair, the number of cCREs and enhancer-like elements within the intervening genomic interval was counted, and mean nucleotide conservation across the path was quantified using phastCons scores derived from a 100-way vertebrate multiple sequence alignment, reflecting purifying selection acting on functional non-coding sequence^56^. Three-dimensional chromatin organisation was incorporated via topologically associating domain (TAD) boundaries and chromatin loop anchors from seven human cell lines (GM12878, HMEC, HUVEC, IMR90, K562, KBM7, NHEK), enabling per-cell-type assessment of whether a variant and its putative target gene reside within a shared TAD or are connected by a chromatin loop. TAD calls were derived from in situ Hi-C data using the directionality index method and loop calls from the HiCCUPS algorithm, both as reported^57^. Finally, recombination hotspot features, including the count of hotspot intervals, peak recombination rate, and a binary indicator of hotspot interposition (>10 cM/Mb), were derived from the HapMap-based fine-scale recombination map accessed via the UCSC Genome Browser recombination rate track. This provided a population-genetic measure of historical recombination intensity that is known to constrain the extent of LD and thereby influence the effective range of regulatory influence^58^.

Across each collated feature, we performed association testing for differences across effector genes contained quasi true-positive sets. Regression models were chosen based on the type of outcome, using linear regression models for continuous variables, logistic regression for binary variables (e.g., variant within a cis regulatory element), and negative binomial regression for counts (e.g., number of cis regulatory elements in the region).

### Analysing GWAS data for six distinct human phenotypes

We obtained lead signals for four European ancestries based GWAS (rheumatoid arthritis, hypothyroidism, prostate cancer, ischaemic heart disease) and GWAS summary statistics for two diseases, namely type 2 diabetes and schizophrenia. Summary statistics were clumped with PLINK2, the minimum p-value for index variants was set to 5×10^-8^, the respective r^2^ of the clump was set to 0.1. The radius from the index variant was set to 500kb for schizophrenia and 1000kb for the type 2 diabetes summary statistics.

### Pathway enrichment

We tested each effector-gene set for over-representation of KEGG^59^ and Reactome^60^ pathways (human gene sets, restricted to 5–500 genes and requiring at least two query genes per set) using a one-sided Fisher’s exact test implemented in a custom R routine. To control for the well-described tendency of highly annotated, “multifunctional” genes to generate biologically non-specific and statistically fragile enrichment^61^, we applied the predictive-specificity correction of Ballouz and colleagues^62^. To this end, within each query, genes were ranked by multifunctionality and the most multifunctional 5% were dropped, and enrichment was recomputed to yield a corrected odds ratio and corrected P value for every term. Multiple testing was controlled across tested sets by the Benjamini-Hochberg false discovery rate (FDR) applied to the corrected P values. A term was considered recovered by a classifier at corrected FDR < 0.05, corrected odds ratio > 1, and at least three overlapping genes. To permit threshold-free comparison between classifiers, we retained every tested term together with its corrected odds ratio and P value rather than only terms passing correction. The correction additionally flagged each term as “specific” (not attributable to generic multifunctional genes) and “robust” (significant before and after gene dropping). We carried these flags through as descriptive annotations to identify potentially redundant, hub-driven enrichments, but did not use them to threshold the signal. The tested background was defined as the union of genes annotated to at least one tested KEGG or Reactome pathway and was identical across classifiers within each source, so between-classifier comparisons were not confounded by differing universes.

### Cell-type enrichment

We tested the same effector-gene sets against cell-type marker sets derived from the Human Protein Atlas single-cell transcriptomic map (v25)^63^. For each of the atlas cell types we defined a marker set as the genes whose expression exceeded the cross-cell-type mean by at least 1.5-fold on the log scale, subject to a minimum expression floor, and tested over-representation of the classifier’s genes among these markers using the same multifunctionality-corrected over-representation test, against a universe of all genes detected in the atlas. Significance was defined at corrected FDR < 0.05 with a corrected odds ratio > 1, and at least three overlapping genes.

#### Redundancy-aware pathway modules

Because pathway gene sets overlapped extensively, comparing classifiers at the level of individual terms would have conflated genuine disagreement with pathway redundancy. We therefore collapsed terms into modules before any comparison. We derived a term-term similarity from the genes shared between terms, pathways or cell type signatures, across the pooled candidate sets using the mean of the Jaccard and overlap coefficients, clustered terms by average-linkage hierarchical clustering on one minus this similarity and cut the dendrogram so that terms sharing at least 30% combined gene overlap formed a single module. Module counts and all downstream metrics were stable across a range of cut heights.

### Cross-classifier consistency

For each disease we quantified agreement among two nearest-gene and five trained classifiers as a threshold-free Spearman rank correlation, across the union of tested modules, of a signed enrichment score defined as the negative log_10_ corrected P value multiplied by the sign of the corrected odds ratio relative to one, so that concordant enrichment or concordant depletion counted as agreement and opposite directions as disagreement. Because this score is continuous in the corrected P value, residual multifunctionality-driven signal were down-weighted smoothly rather than by a hard filter, keeping the score comparable across classifiers. Overall agreement per disease was summarised as the mean off-diagonal similarity of the resulting classifier-by-classifier matrices.

### Classifier distinctiveness and its stability across diseases

To identify classifiers that systematically recovered distinct biology, we counted, per disease and classifier, the modules recovered exclusively by that classifier (recurrence of one) and its mean agreement with the remaining classifiers. We then tested whether the same classifier was consistently the most distinct across diseases by applying Kendall’s coefficient of concordance (with the associated Friedman test) to the disease-by-classifier matrix of mean agreement. We used the mean rank to identify the systematically deviant classifier.

#### Effect sizes and gene annotation

Multifunctionality-corrected odds ratios were the primary effect measure for both pathways and cell types, with the representative (“driver”) term of each module defined as its most significantly enriched member. Gene sets and effector genes were harmonised on HGNC gene symbols, with Ensembl identifiers mapped to symbols using the Ensembl GRCh38 gene annotation, release 112^64^.

### Drug-target recovery benchmark

We curated a second dataset of GWAS loci that recapitulate known drug–target–indication triplets. To this end, we first compiled a systematic list of approved or Phase IV therapeutic targets and their associated indications from the Open Targets Platform (v.25.07), and then collected GWAS from an updated release of the GWAS Catalog for the indications with such approved drug targets. This mapping yielded an initial set of 2,705 genome-wide significant loci (p < 5×10⁻⁸) with an approved drug target and indication (i.e. triplets). In contrast to the benchmarking dataset, no manual review was performed. Finally, we defined 250 kb genetic regions to collapse overlapping signals for the same indication–drug target pair, resulting in 340 unique locus–indication–drug target triplets. (**Supplementary Fig. 5**).

Classifiers were evaluated for recovery of established drug targets across 340 locus-target-indication triplets in which the approved target lay within 1 Mb of the lead variant for the respective indication. Candidate-gene scores were min-max scaled within each locus, and recovery was summarised per classifier as the within-locus area under the ranking curve (the probability that the target outranks a randomly chosen competing gene at the locus) together with the proportion of loci at which the target ranked first (rec@1) or within the top three (rec@3). Differences in recovery rate across classifiers were tested by Cochran’s Q test on the binary rec@1 matrix and by the Friedman test and Kendall’s W on the within-locus AUC matrix. For targets recovered by a single classifier, the recovering model’s score margin over the next-best candidate was compared with the target’s median rank under the remaining classifiers to distinguish credible recoveries from single-model nominations against cross-model consensus.

### Benchmark against Open Targets L2G

We compared DEFA with Open Targets locus-to-gene^13^ (L2G) on the Open Targets 26.06 all-disease export, taking one credible set-trait pair as the unit (180,017 pairs) and restricting all comparisons to the 178,518 pairs for which every method made a call. We evaluated three DEFA model sets: knowledge-free (the five domain-informed classifiers without knowledge-graph features), knowledge-informed (the same classifiers with graph features) and label-free (the knowledge-free set without the Open Targets-informed classifier, whose training labels include drug targets). Per set we assigned the gene with the highest probability across classifiers (primary), with the lowest sum of median and maximum probability ranks, or with the highest mean probability (sensitivities). Reference methods were the L2G top gene, the nearest protein-coding gene by TSS and the closest gene by regulatory footprint.

We scored top-1 accuracy against three lines of evidence, each on pairs where at least one candidate gene carried the evidence: rare-variant gene-burden associations for the trait [burden source; 5,737 pairs]; Mendelian gene–disease relations from Orphanet, Gene2Phenotype, ClinGen and Genomics England (2,000 pairs); and an approved drug for the gene in the same disease according to Open Targets clinical precedence (4,500 pairs). Because L2G was trained on curated gene-locus pairs that include drug targets, we matched its training file to the 26.06 credible sets by credible set, else by lead variant (83.8% of positives placeable), and defined pairs outside L2G training as those whose credible set was not a training positive and that carried neither of two leakage flags: (A) the pair’s gene-disease group or genomic locus contained a training-positive credible set, or (B) a true gene was a training positive at a credible set sharing a variant or with a lead variant within 500 kb, for any trait. These flags are lower bounds (**Supplementary Tab. 10**). The approved-drug test was evaluated primarily on the 770 pairs outside L2G training, the Mendelian test on all 2,000 pairs and on the 660 outside L2G training.

Because the same gene–disease relation recurs across studies and overlapping credible sets, we grouped pairs sharing any (true gene, evidence disease) relation into linked groups (540, 277 and 339 for the three primary frames; largest group 3.6% of pairs, 5.5% for *CFHR1-4* in the Mendelian frame on non-training credible sets; **Supplementary Tab. 10**) and derived all confidence intervals from a paired bootstrap over groups (2,000 resamples). Grouping by gene or by exact gene set gave the same intervals; weighting pairs by the inverse group size or keeping one pair per group (the credible set with the smallest lead P) are reported in **Supplementary Tab. 10**.

To locate the source of the knowledge-informed gain, we ranked each pair’s candidate genes by single graph features (shortest-path distance, personalised PageRank, Adamic–Adar, resource allocation and degree in PrimeKG^15^, release 2022-04-25) and stratified pairs, before computing any result, by whether a true gene was the unique candidate at the shortest distance to the trait, tied with others, or not the closest. Within strata we contrasted knowledge-informed with knowledge-free DEFA and with L2G, and estimated the difference-in-differences between strata from the same group resamples; PageRank as stratifier served as sensitivity. As the graph’s sources predate July 2021, we repeated the comparison with L2G on pairs whose true genes were all first reported after 2021 (2022 and 2023 as sensitivities).

### Data analysis

All analyses and most image generation were performed in Python (v3.12) or R (v4.4.1). In Python, we used polars (v1.40), pandas (v2.3.3), numpy (v2.3.5), scipy (v1.16), pyarrow (v23), and matplotlib (v3.10). In R, we used data.table (v1.17), ggplot2 (v4), and dplyr (v1.1.4).

**Supplementary Note 1 | Classifier-private disease biology**

Ischaemic heart disease. Genes with enhanced expression in cardiomyocytes, uniquely enriched in the mQTL-informed model (OR=1.97; FDR=4.5×10-2), comprised canonical coronary risk genes *ADAMTS7*, *PHACTR1*, *EDNRA*, *LPL*, and *PLPP3* alongside genes with established roles in cardiomyocyte identity, sarcomere composition, and cardiac development, like the cardiac transcription factor *TBX20*, the cardiomyopathy gene *PRDM16*, and the sarcomeric myosin *MYH7B*^24^. For the complement and coagulation cascade, recovered only by the FLAMES-informed classifier, interventional evidence is confined to the coagulation arm: in the randomised COMPASS trial, adding the direct factor Xa inhibitor rivaroxaban to aspirin reduced a composite of cardiovascular death, stroke or myocardial infarction by about 24%^25^. Evidence for the complement arm remains preclinical: terminal complement products are found in human atherosclerotic plaques at higher levels in unstable angina and myocardial infarction than in stable lesions^26^, suppression of C3 activation slowed progression of atherosclerosis in Ldlr-deficient mice^27^, inhibition of the terminal component C5 reduced infarct size and cardiomyocyte apoptosis in myocardial ischaemia-reperfusion^28,29^, and C1-inhibitor, a serine protease inhibitor regulating the complement, coagulation and kallikrein-kinin systems simultaneously, has been proposed as a therapeutic candidate without definitive trial evidence in acute myocardial infarction^30^. Two further examples were private to distinct classifiers: vitamin K-dependent γ-carboxylation (F7, GGCX, and FURIN; pQTL-informed; OR=28.4; FDR=1.5×10-2) and organic cation/anion/zwitterion transport (*SLC22A1* and *SLC22A3-5*; mQTL-informed; OR=15.2; FDR=1.8×10^-2^).

Type 2 diabetes. PPAR signalling illustrates the translational relevance of consistently recovered pathways: PPARγ, encoded by PPARG, is the target of thiazolidinediones and PPARα, encoded by PPARA, of fibrates — ligand-activated nuclear receptors controlling adipocyte lipid storage and insulin sensitisation, and hepatic fatty-acid oxidation and lipoprotein metabolism, respectively^22^. Saturation mutagenesis of all 9,595 possible missense variants in PPARG further established that loss-of-function alleles cause Mendelian lipodystrophy and increase type 2 diabetes risk^23^. Downstream pathway members assigned across classifiers included ADIPOQ, ANGPTL4, LPL, APOA1, APOC3, ACSL1, ACOX2, and CPT1B. Schizophrenia. GRIN2A, central to the long-term potentiation signal recovered only by the regulatory-footprint baseline, carries convergent common- and rare-variant evidence^21,31^, and heterozygous Grin2a loss of function reproduces both hypo-glutamatergic and hyper-dopaminergic features in mice^65^.

Rheumatoid arthritis. CTLA4, within the cell adhesion molecules pathway recovered only by nearest-gene assignment by transcription start site, encodes the T-cell co-inhibitory receptor whose extracellular domain, fused to an immunoglobulin Fc fragment, forms the costimulation modulator abatacept, which improved clinical response in rheumatoid arthritis refractory to TNF inhibition^32^.

