## Supplementary Figures 1-6 for "Domain-informed effector gene assignment exposes training-data dependence of drug target prioritisation from GWAS"

**Supplementary Figure 1 | Performance of published effector gene prediction models on three distinct benchmarking datasets.**

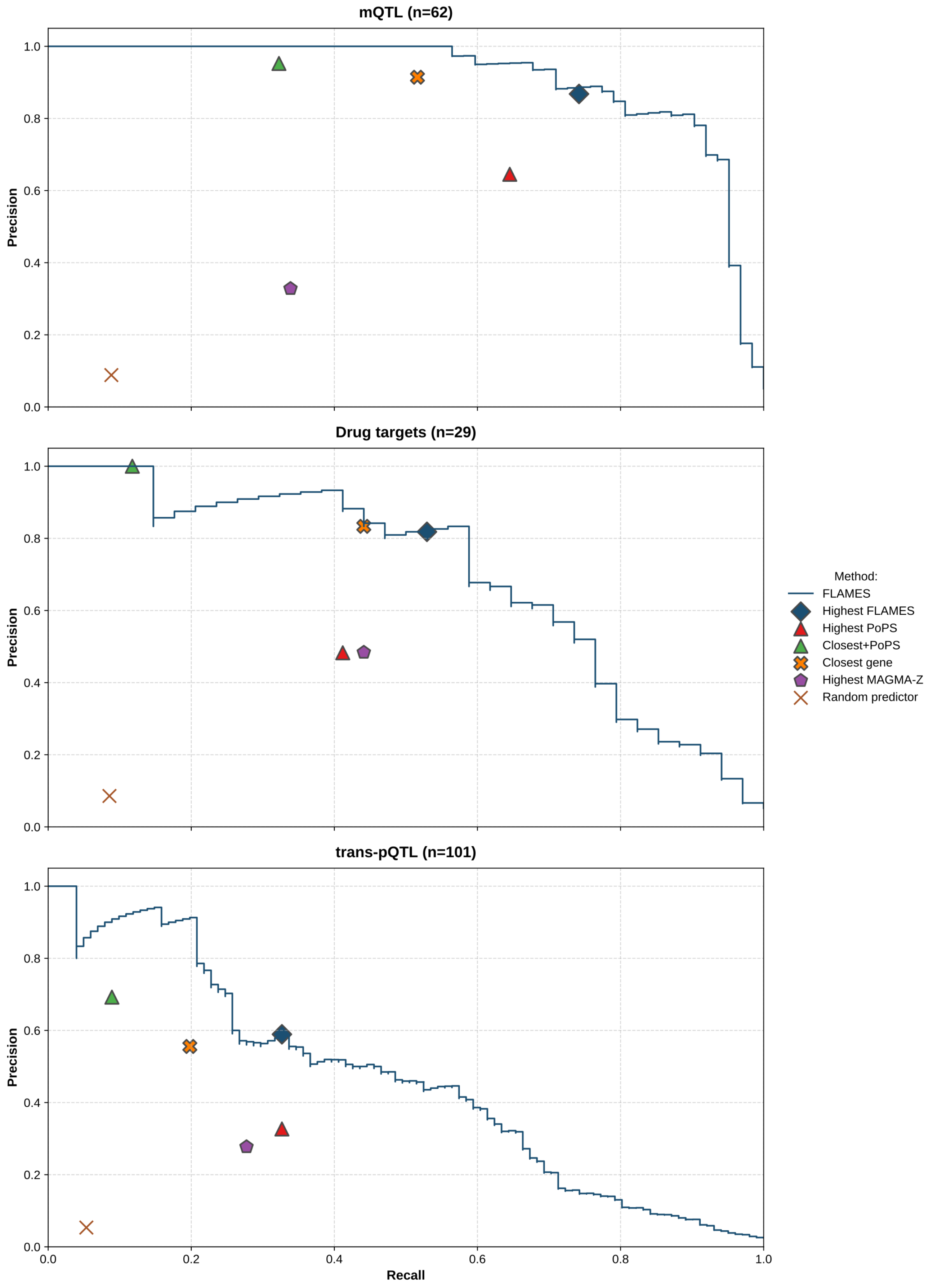

#### Supplementary Figure 2 | Performance of DEFA models on different dataset with confidence-intervals and graph-feature contributions.

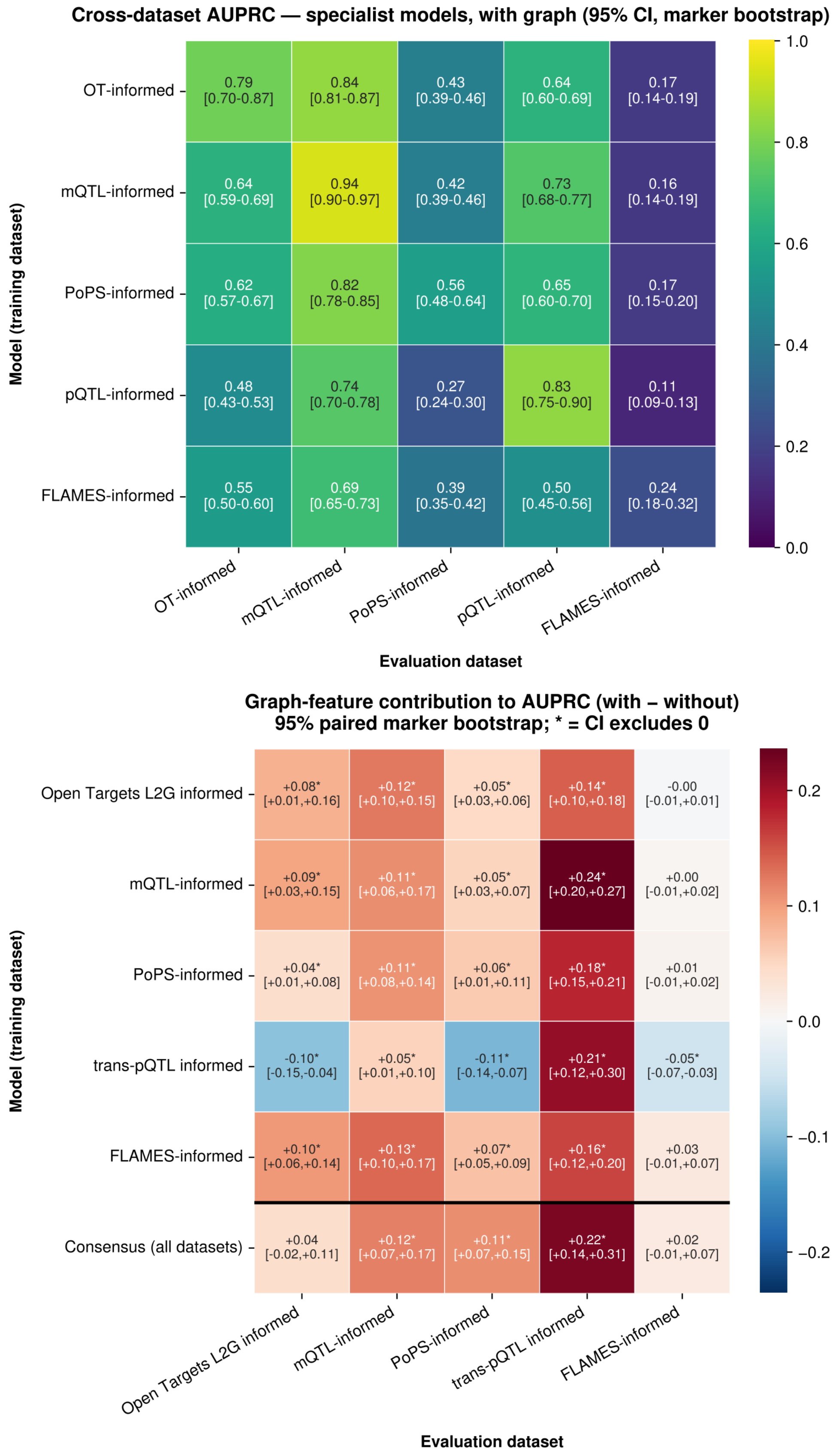

**Supplementary Figure 3 | Evolutionary constraint of genes inside the domain-informed model datasets**

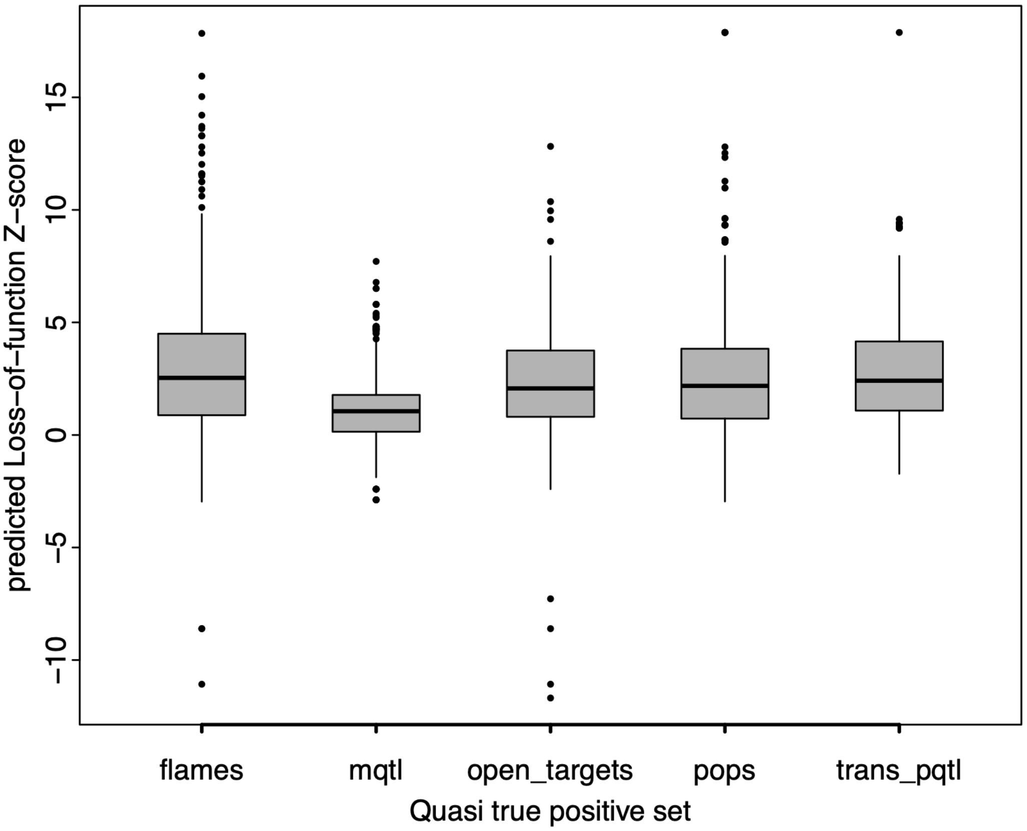

### Supplementary Figure 4 | Sensitivity to the LD proxy r2 threshold

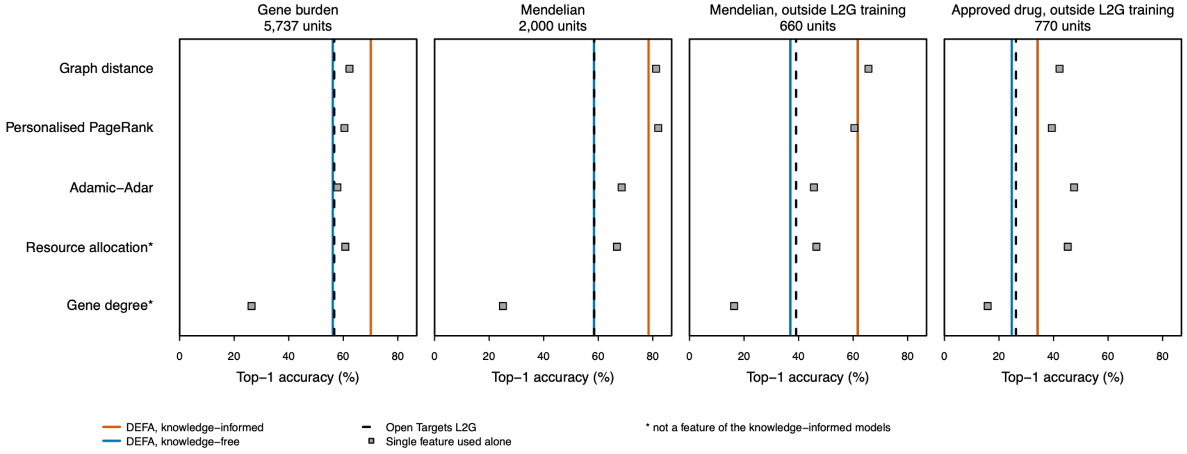

**Supplementary Figure 5 | Creating a dataset of unique Locus-Target -Indication Triplets to estimate the recovery of targets from FDA-approved drugs.**

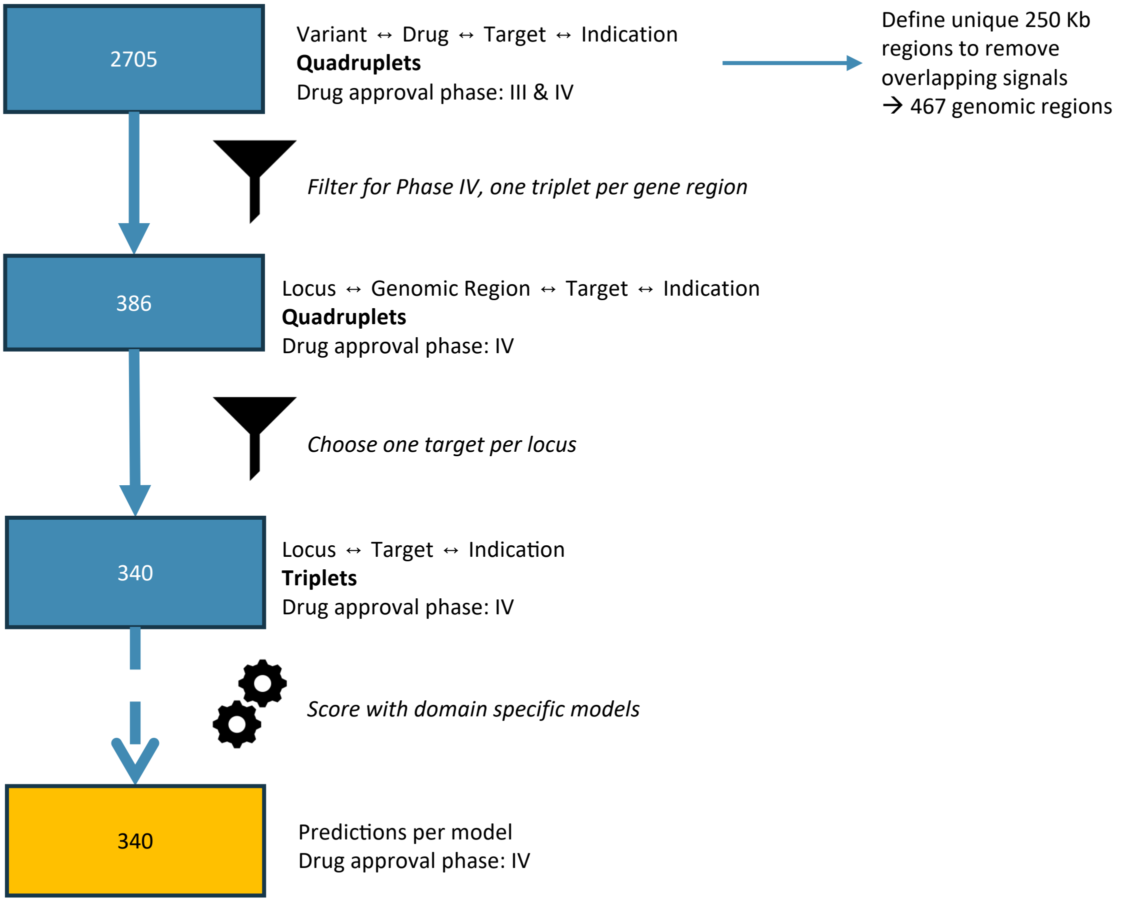

### Supplementary Figure 6 | Sensitivity to the LD proxy r2 threshold

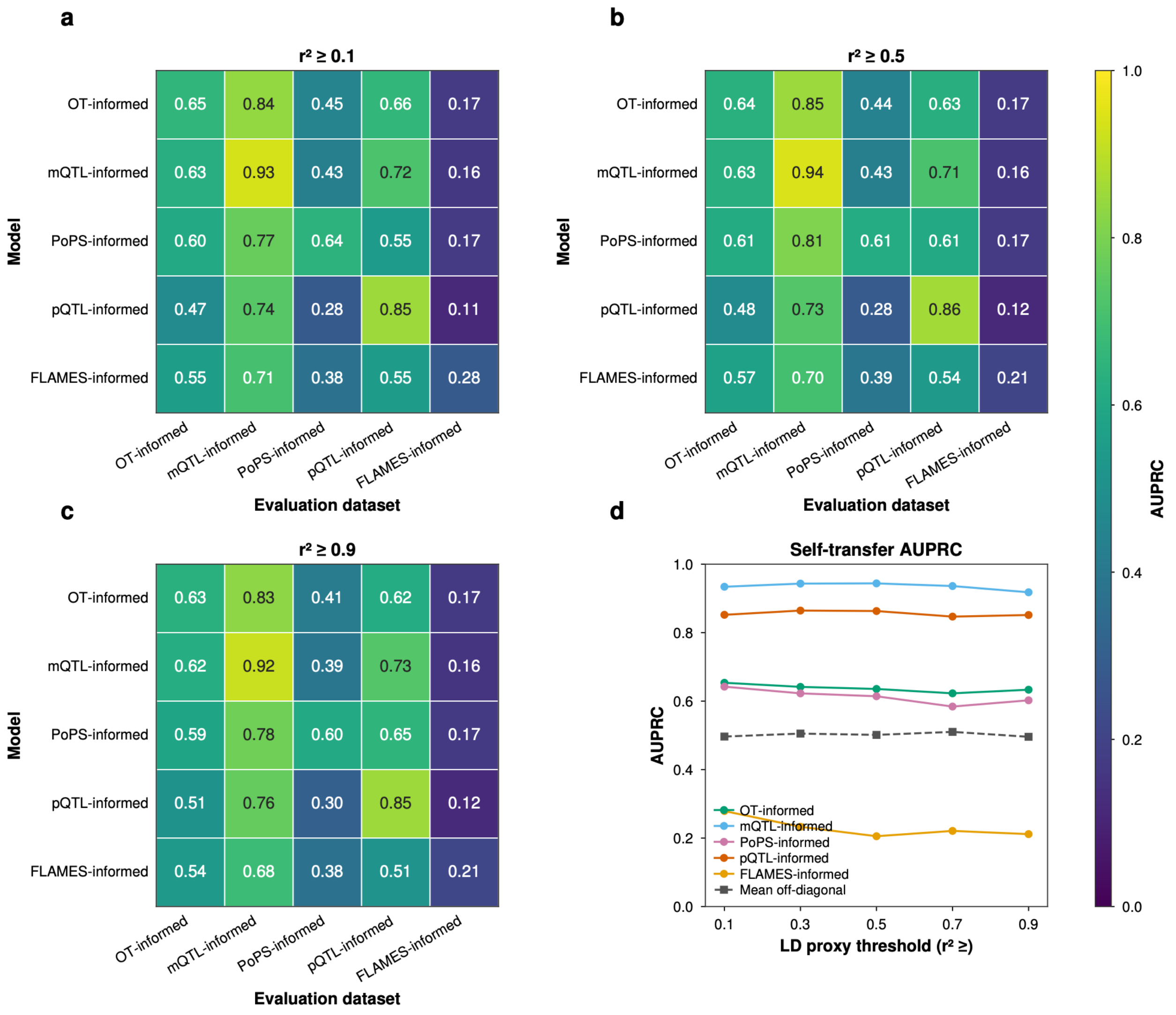
